# Discovery of a molecular glue promoting CDK12-DDB1 interaction to trigger Cyclin K degradation

**DOI:** 10.1101/2020.06.10.144303

**Authors:** Lu Lv, Peihao Chen, Longzhi Cao, Yamei Li, Zhi Zeng, Yue Cui, Qingcui Wu, Jiaojiao Li, Jian-Hua Wang, Meng-Qiu Dong, Xiangbing Qi, Ting Han

**Affiliations:** College of Life Sciences, Beijing Normal University, Beijing, China; National Institute of Biological Sciences, Beijing, China; School of Life Sciences, Peking University, Beijing, China; Graduate School of Peking Union Medical College and Chinese Academy of Medical Sciences, Beijing, China; Tsinghua Institute of Multidisciplinary Biomedical Research, Tsinghua University, Beijing, China

## Abstract

Molecular glues are small molecules that exert their biologic or therapeutic activities by inducing gain-of-function interactions between pairs of proteins. In particular, molecular-glue degraders, which mediate interactions between target proteins and components of the ubiquitin proteasome system to cause targeted protein degradation, hold great promise as a unique modality for therapeutic targeting of proteins that are currently intractable. Here, we report a new molecular glue HQ461 discovered by high-throughput screening of small molecules that inhibited NRF2 activity. Using unbiased loss-of-function and gain-of-function genetic screening followed by biochemical reconstitution, we show that HQ461 acts by promoting interaction between CDK12 and DDB1-CUL4-RBX1 E3 ubiquitin ligase, leading to polyubiquitination and proteasomal degradation of CDK12’s interacting protein Cyclin K (CCNK). Degradation of CCNK mediated by HQ461 compromised CDK12 function, leading to reduced phosphorylation of CDK12 substrate, downregulation of DNA damage response genes, and cell death. Structure-activity relationship analysis of HQ461 revealed the importance of a 5-methylthiazol-2-amine pharmacophore and resulted in an HQ461 derivate with improved potency. Our studies reveal a new molecular glue that engages its target protein directly with DDB1 to bypass the requirement of a substrate-specific receptor, presenting a new strategy for targeted protein degradation.

## Introduction

Molecular glues are a class of small molecules that induce the formation of protein-protein interactions to elicit biologic or therapeutic effects. The name “molecular glue” was first coined to describe the mechanism of action of the plant hormone auxin, which bridges an interaction between the E3 ubiquitin ligase TIR1 and IAA transcription repressors, leading to IAA destruction by the ubiquitin proteasome system to activate auxin-response gene expression^1^. Unlike conventional small molecules that bind to active sites or allosteric sites to modulate the activity of their target proteins, molecular glues influence the activity or fate of their target proteins by bringing them to the vicinity of a regulatory protein. As a result, molecular glues have been viewed enthusiastically as a unique pharmacological modality to target proteins without druggable pockets^2^.

Notwithstanding, molecular glues are rare and were discovered serendipitously; only a handful of molecular glues have been documented over the past four decades. For example, macrocyclic natural products cyclosporin A, FK506, and rapamycin recruit FKBP proteins to the phosphatase calcineurin or the kinase mTORC1, interfering with their enzymatic activities to control intracellular signal transduction^3,4^. Similarly, the thalidomide class of immunomodulatory drugs binds to the E3 ubiquitin ligase cereblon and alters its substrate specificity to recognize and degrade several zinc finger transcription factors^5–7^. More recently, indisulam and related anti-cancer sulfonamides were discovered to function by promoting the interaction between the splicing factor RBM39 and the E3 ubiquitin ligase DCAF15, resulting in the degradation of RBM39 to cause aberrant pre-mRNA splicing^8,9^.

Several common themes have emerged from this limited set of molecular glues. First, molecular glues typically bind to their target proteins with modest or even undetectable affinities^8^; while enhanced affinities are usually observed once a regulatory protein is also present in the system. Such unique binding characteristics can be explained by induced protein-protein interactions and pre-existing protein structural complementarity as revealed by structural analysis of molecular glue-bound protein complexes^1,10–12^. Second, molecular glues can target transcription factors and splicing factors, by recognizing either relatively flat surfaces or disordered regions of these proteins^6–8^. Third, molecular glues possess favorable pharmacological properties to serve as drug candidates. For example, the compactness of the thalidomide scaffold allowed the development of bivalent proteolysis target chimeras to direct cereblon to degrade disease-relevant proteins^13,14^. However, the rarity of molecular glues has limited their potential as a general strategy for drug development.

In this study, we report the discovery and the mechanism of action of a new molecular glue HQ461. HQ461 was identified through phenotype-based high-throughput small-molecule screening and found to possess potent cytotoxicity. Combining chemical genetics and biochemical reconstitution, we show that HQ461 acts by binding to the ATP-binding pocket of CDK12’s kinase domain, creating a binding interface for DDB1, which is a subunit of the DDB1-CUL4-RBX1 E3 ubiquitin ligase. Distinct from existing molecular glues that engage the ubiquitin proteasome system by binding to a substrate-specific receptor, HQ461 converts CDK12 into a substrate-specific receptor to trigger the polyubiquitination and subsequent degradation of CDK12’s partner protein Cyclin K (CCNK). HQ461-mediated depletion of CCNK from cells led to reduced Serine 2 phosphorylation of RNA polymerase II CTD and affects the expression of genes involved in DNA damage responses.

## Results

### Discovery of HQ461 by high-throughput screening

NRF2 is a transcription factor that is polyubiquitinated by the CUL3-KEAP1 ubiquitin ligase and degraded by the proteasome under normal conditions^15^. Oxidative stress or xenobiotic exposure inactivates KEAP1 to stabilize NRF2, which subsequently activates expression of genes related to antioxidant response and xenobiotic detoxification^15^. Clinically, *KEAP1* mutations are frequently found in a variety of cancers^16,17^, raising the possibility that inhibiting overactive NRF2 may provide therapeutic benefits to patients with *KEAP1* mutations. To identify small-molecule NRF2 inhibitors, we used an NRF2 activity-based luciferase reporter (ARE-luc2P) assay in the lung adenocarcinoma cell line A549, which expresses high levels of NRF2 due to a *KEAP1* mutation (Fig. 1A) and performed high-throughput screening of a library containing 65,790 small molecules at a concentration of 10 μM. After Z-score normalization, 515 compounds (Z-score < −2.5, hit rate 0.78%) were found to reduce ARE-*luc2P* activity (Fig. S1A).

**Figure 1.**
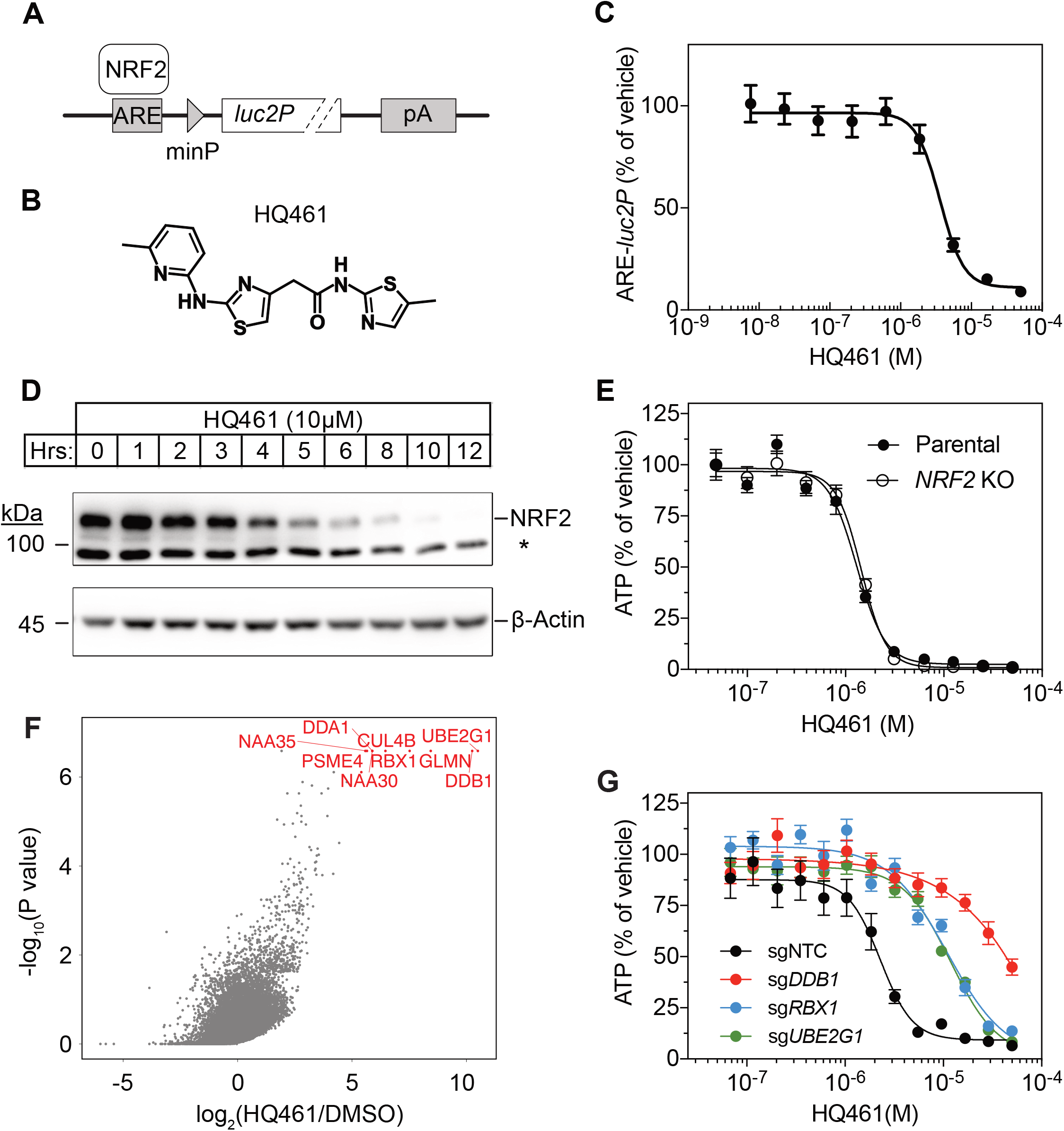
DDB1-CUL4-RBX1 mediates HQ461’s cytotoxicity. (A) ARE-*luc2P* reporter used for high-throughput screening. NRF2 binds to anti-oxidant response elements (ARE) to drive luciferase (*luc2P*) expression. (B) Chemical structure of HQ461. (C) Measurement of the HQ461 IC_50_ on ARE-*luc2P* activity. Error bars represent standard errors of the mean (SEM) from three biological replicates. (D) HQ461 treatment led to reduced NRF2 protein level in A549 cells. The asterisk indicates a non-specific band. (E) Measurement of the HQ461 IC_50_ on the viability of *NRF2*-knockout and parental A549 cells. Error bars represent SEM from three biological replicates. (F) MAGeCK analysis of pooled genome-wide CRISPR-Cas9 sgRNA screening of HQ461 resistance in A549 cells. (G) Measurement of the HQ461 IC_50_ on the viability of A549 cells expressing non-targeting control (NTC) or sgRNAs targeting *DDB1*, *RBX1*, or *UBE2G1*. Error bars represent SEM from three biological replicates.

To exclude general toxins that interfere with global gene expression and inhibitors of luciferase enzymatic activity, we counter-screened these 515 compounds by measuring their activities towards a luciferase reporter driven by the herpes simplex virus thymidine kinase promoter (TK-*luc2P*). Fifty-six compounds exhibited greater than 2.5-fold selectivity towards ARE-*luc2P* relative to TK-*luc2P* (Fig. S1B). For this paper, we focused on HQ461 (Fig. 1B), which features an aminopyridinylthiazol scaffold (Fig. 1B); this compound inhibited ARE-*luc2P* activity at a half maximal inhibitory concentration (IC_50_) of 3.6 μM (Fig. 1C). By western blotting and RT-qPCR, we found that HQ461 treatment led to a rapid reduction of NRF2 mRNA and protein levels (Fig. 1D and S1C). These results suggest that HQ461 functions upstream of NRF2 transcription. In addition to an effect on NRF2 expression, HQ461 displayed potent cytotoxicity in A549 cells (IC_50_ of 1.3 μM). To examine if NRF2 downregulation may be the cause of HQ461’s cytotoxicity, we used CRISPR-Cas9 to inactivate *NRF2* in A549 cells and were able to isolate an NRF2 knockout clone, which was equally sensitive to HQ461’s cytotoxicity relative to its parental cell line (Fig. 1E). These observations suggest that HQ461’s cytotoxicity is independent of NRF2.

### HQ461’s cytoxicity requires DDB1-CUL4-RBX1

To investigate HQ461’s mechanism(s) of action, we performed pooled genome-wide CRISPR-Cas9 knockout screening by targeting 19,114 genes with four individual sgRNAs per gene^18^. We treated sgRNA-transduced A549 cells with either the vehicle DMSO or a sub-lethal dose of HQ461 for three weeks to enrich for cells that were resistant to HQ461’s cytotoxic effects. Afterwards, we isolated genomic DNAs from surviving cells and performed sgRNA amplification by PCR followed by next generation sequencing to measure the abundance of each sgRNA in these cells. Using the MAGeCK algorithm^19^ to rank enriched genes in HQ461 treated relative to DMSO treated cells, we found the top-ranking genes include the E2 ubiquitin activating enzyme G1 (*UBE2G1*), three members of the DDB1-CUL4-RBX1 E3 ubiquitin ligase complex (*DDB1*, *RBX1*, *DDA1*, *CUL4B*), and multiple regulators of DDB1-CUL4-RBX1 activity (*GLMN*, *NAA30*/*35*/*38*, *NAE1*, and *UBE2M*) (Fig. 1F, S1D, and Table S1). We targeted three top-ranking genes *DDB1*, *RBX1*, and *UBE2G1* individually by two independent sgRNAs; expression of these sgRNAs in A549 cells resulted in the depletion of their target proteins (Fig. S1E) and resistance to HQ461’s toxicity (Fig. 1G and S1F). Because all these candidate genes encode proteins in the ubiquitin proteasome system, we hypothesized that HQ461 may exert its cytotoxicity by triggering proteasomal degradation of target protein(s).

### *CDK12* mutations cause HQ461 resistance

To identify any target protein(s) destabilized by HQ461, we performed gain-of-function genetic screening in the colorectal cancer cell line HCT-116. HCT-116 cells are defective in mismatch repair, and therefore harbor a high rate of random point mutations^20,21^. Previous studies suggest that the identification of recurrent mutations present in multiple independent drug-resistant clones may reveal the direct drug target^8,21^. To ensure the isolation of independent HQ461-resistant clones, we derived 10 clonal isolates from HCT-116 that were initially sensitive to HQ461 (HQ461^S^). Five of these clones evolved HQ461 resistance (HQ461^R^) during the expansion from a single cell into a large cell population (Fig. 2A). We isolated one HQ461^R^ clone from each cell population and confirmed their resistance to HQ461 (3-fold increase of IC_50_ relative to parental HCT-116 cells) (Fig. 2B). We pooled genomic DNAs isolated from the five HQ461^R^ clones at equal amounts for whole-exome sequencing. As a control, we sequenced the exomes of their corresponding HQ461^S^ clones as a pool.

**Figure 2.**
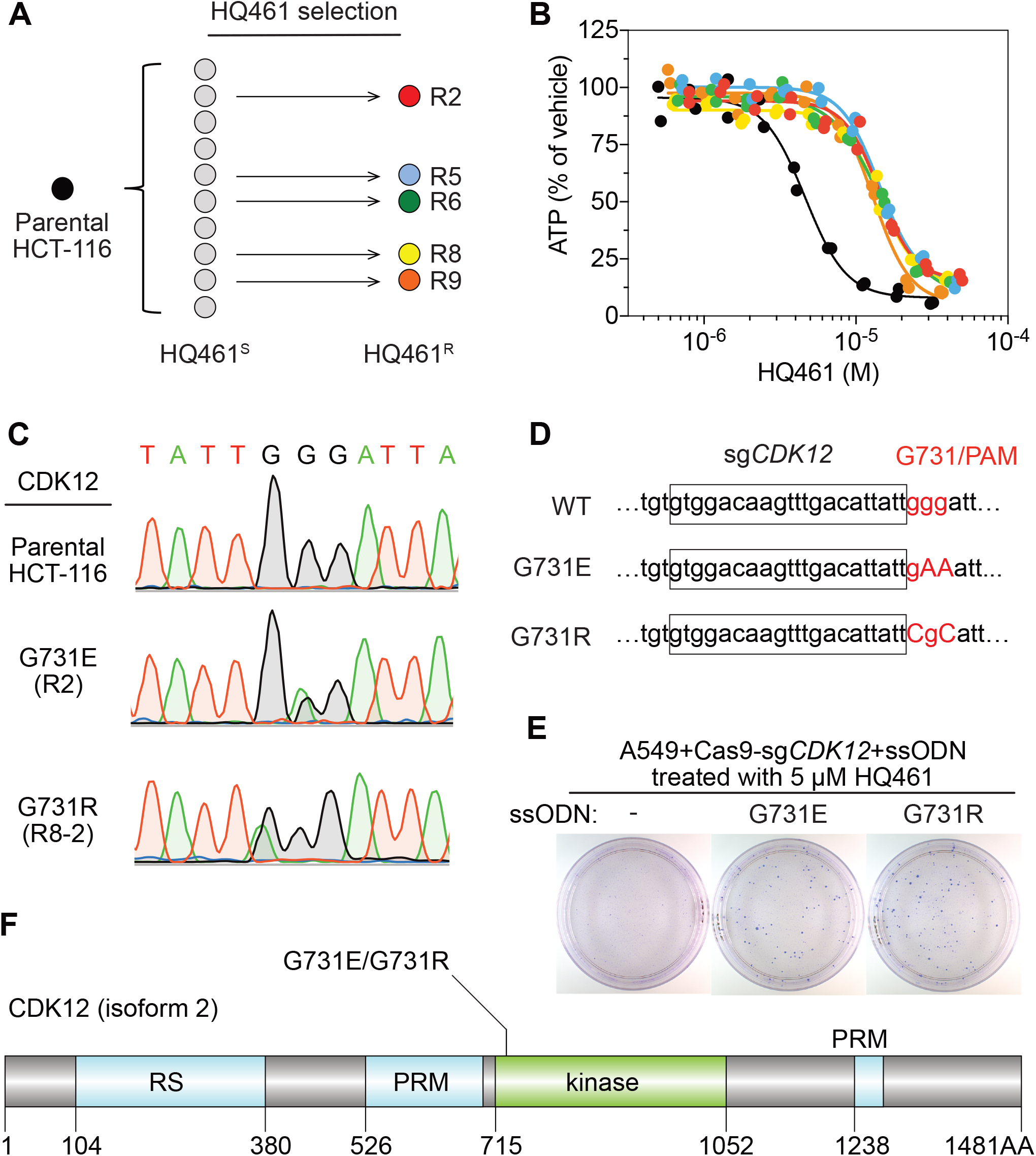
Mutations in CDK12 confer resistance to HQ461. (A) Strategy for isolation of independent HQ461-resistant clones. (B) Measurement of the HQ461 IC_50_ on the viability of parental and five HQ461 resistant clones of the A549 cell line (color coded in (A)). Error bars represent SEM from three biological replicates. (C) Sanger sequencing verification of *CDK12* mutations found in HQ461 resistant cells. (D) The edited genomic sequence of *CDK12*, highlighting the PAM motif, the target sequence (boxed), and G731E or G731R mutations. (E) *CDK12* G731E or G731R knock-in in A549 cells gives rise to HQ461 resistance, visualized by crystal violet staining. Mock transfection omitting the repair template results in no HQ461 resistance. (F) The domain structure of CDK12 (isoform 2 is shown here and used in this study). G731 is located within the central kinase domain of CDK12.

We then used the GATK pipeline^22–24^ to identify variants that were unique to the HQ461^R^ group (Table S2). The top-ranking variant was a G to A substitution resulting in a non-synonymous substitution of glycine to glutamate (G731E) in the gene Cyclin-dependent kinase 12 (*CDK12*). This variant occurred at an allele frequency of ~30% in HQ461^R^ versus 0% in HQ461^S^ (Table S2). Deconvolution of pooled samples by Sanger sequencing of *CDK12* revealed a heterozygous *CDK12* G731E mutation in three out of five clones, consistent with an overall allele frequency of 30% in a diploid cell line (Fig. 2C). Using Sanger sequencing of *CDK12* to examine additional HQ461^R^ clones, we discovered another *CDK12* mutation affecting the same codon which replaces glycine with arginine (G731R) (Fig. 2C).

To test if G731E and G731R mutations in *CDK12* confer HQ461 resistance, we knocked-in these two mutations in A549 cells using CRISPR-Cas9 enhanced homology dependent repair. We designed a sgRNA targeting a 21-nucleotide sequence upstream of codon G731 of *CDK12*. G731 is encoded by the codon GGG, serving conveniently as a protospacer adjacent motif (PAM) for Cas9^25^. To distinguish knock-in mutations from spontaneous mutations, we designed two-nucleotide substitutions on the repair templates to encode G731E (GGG to GAA) or G731R (GGG to CGC) (Fig. 2D). Successful editing of GGG sequence to GAA or CGC destroys the PAM sequence, preventing the Cas9/sgRNA complex from re-cutting.

We co-transfected plasmids expressing Cas9, the *CDK12* targeting sgRNA, and single-stranded oligodeoxynucleotides (ssODNs) encoding the G731E or G731R allele. These conditions increased clonal resistance to HQ461 (Fig. 2E). Four independent clones were isolated. They contained the expected G731E or G731R mutation (Fig. S2A) and were between 9 to 13-fold less sensitive to HQ461 than parental A549 cells (Fig. S2B). Extending from these observations, we generated stable cell lines expressing either wild-type CDK12 or CDK12 carrying G731E or G731R substitution. Whereas wild-type CDK12 did not alter A549’s sensitivity to HQ461, expression of CDK12 G731E or G731R resulted in HQ461 resistance (Fig. S2C).

### HQ461 triggers polyubiquitination and proteasomal degradation of Cyclin K

CDK12 is a large protein with a central kinase domain flanked by an arginine serine rich (RS) domain and two proline rich motifs. The G731 hotspot for HQ461 resistance mutations is located in the kinase domain of CDK12 (Fig. 1F). This structural position for the mutation led us to speculate that CDK12 might be a protein substrate destabilized by DDB1-CUL4-RBX1 following HQ461 treatment and that G731E or G731R substitution in CDK12 might interfere with such destabilization. By monitoring the protein level of CDK12 at different time points of HQ461 treatment, we found a modest 50% reduction of CDK12 protein level after 8 hours of treatment.

To examine if a 50% reduction of CDK12 protein might help explain HQ461’s cytotoxicity, we used CRISPR interference (CRISPRi)^26^ to downregulate *CDK12* transcription. We used ten independent sgRNAs targeting the transcription start site of *CDK12* and found all of them efficiently depleted CDK12 mRNA and protein (between 5 to 10-fold reduction) without causing cell death or a proliferation defect (Fig. S3A). This result suggested that modest CDK12 downregulation by HQ461 is not a direct cause of cell death. CDK12 binds to Cyclin K (CCNK) to form a functional and active complex to perform its functions^27,28^. In contrast to the modest effect on CDK12 protein, HQ461 treatment led to > 8-fold reduction of the CCNK protein level after a short 4-hour treatment (Fig. 3A). In cells with CDK12 G731E or G731R mutation, CCNK protein was not affected by HQ461 treatment (Fig. 3B).

**Figure 3.**
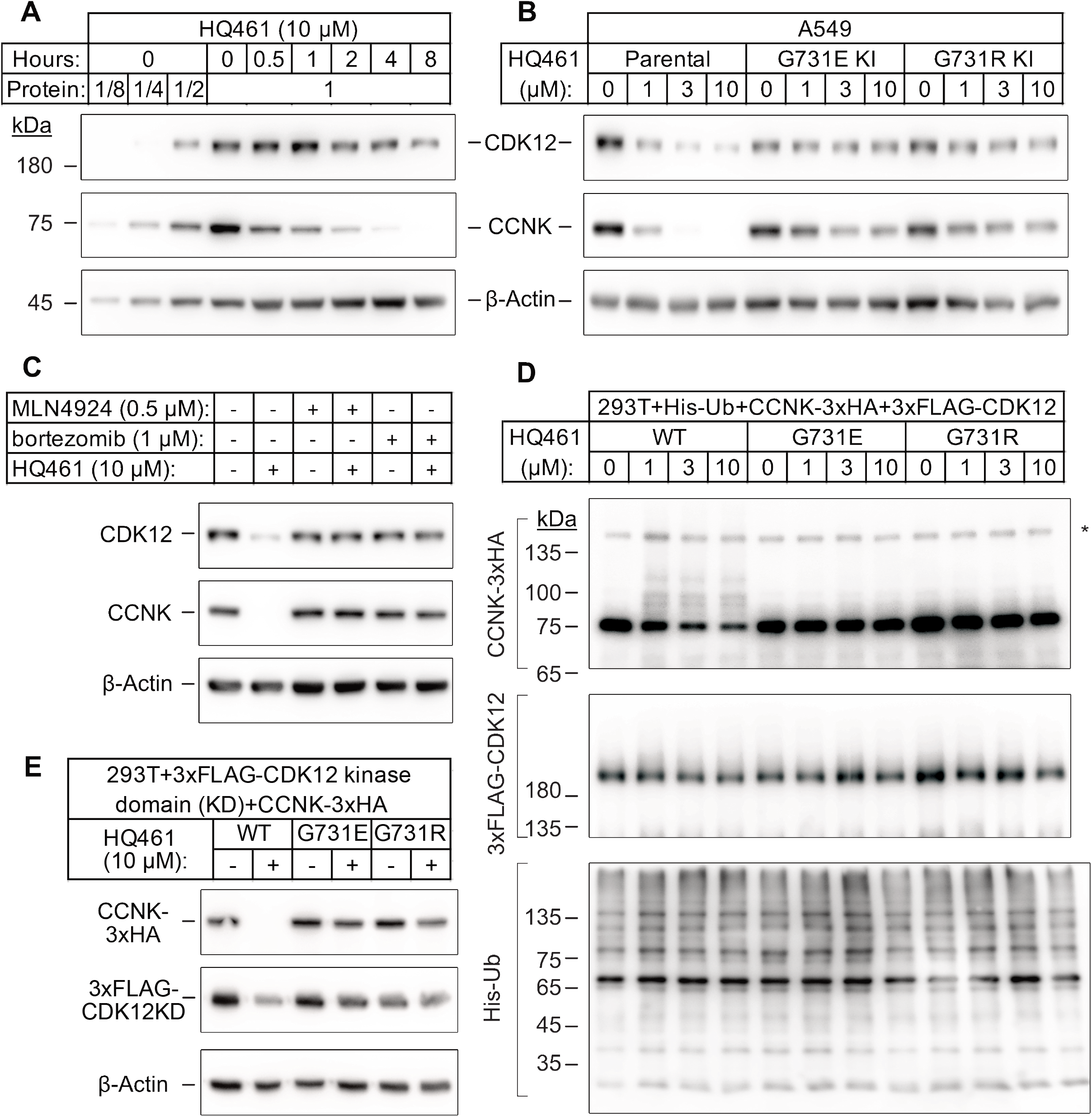
HQ461 promotes CCNK polyubiquitination and proteasomal degradation through DDB1-CUL4-RBX1. (A) Western blotting of CDK12 and CCNK from cells treated with HQ461 treatment. Dilutions of untreated sample (0-hour) were included to allow quantitative assessment of the reduction of protein levels. (B) Effect of *CDK12* G731E and G731R mutations on CCNK and CDK12 protein levels in A549 cells treated with HQ461. (C) Effect of bortezomib and MLN4924 on CCNK and CDK12 protein levels in A549 cells treated with HQ461. (D) HQ461 triggers in vivo polyubiquitination of CCNK in complex with wild-type CDK12 but not CDK12 G731E or G731R. (E) Wild-type CDK12 kinase domain but not G731E or G731R mutants is sufficient for HQ461-mediated destabilization of CCNK.

To test if CCNK depletion requires a functional ubiquitin proteasomal system, we used a proteasome inhibitor bortezomib^29^ and a neddylation inhibitor MLN4924^30^ (inactivating all Cullin RING ligases including DDB1-CUL4-RBX1) and found both inhibitors abrogated HQ461-dependent CCNK degradation (Fig. 3C). Furthermore, in cells depleted of DDB1 or RBX1, CCNK remained stable following HQ461 treatment (Fig. S3B). We next sought to examine if HQ461 triggered CCNK polyubiquitination, so we co-transfected HEK293T cells with plasmids encoding 6xHis-tagged ubiquitin, 3xHA-tagged CCNK, and either wild-type 3xFLAG-tagged CDK12 or mutants with G731E/R substitution. After purification of lysates by nickel chromatography, we performed western blotting with anti-HA and anti-FLAG antibodies to examine if CDK12 and CCNK were polyubiquitinated in cells treated with HQ461. We observed increased CCNK polyubiquitination following HQ461 treatment when wild-type CDK12 was co-expressed. In contrast, co-expression of CDK12 G731E or G731R impeded CCNK polyubiquitination. Furthermore, neither wild-type or mutant CDK12 was polyubiquitinated. Taken together, these results support that HQ461 triggers CCNK polyubiquitination as mediated by DDB1-CUL4-RBX1 and that G731E/R mutations in CDK12 prevent CCNK polyubiquitination.

CDK12 is known to interact with CCNK via its kinase domain^31,32^, and this domain was sufficient to mediate both CCNK polyubiquitination and degradation (Fig. 3E and S3C). Similar to full-length CDK12, co-expression of CDK12’s kinase domain with G731E/R mutations blocked CCNK polyubiquitination and degradation (Fig. 3E and S3C). CDK13 is a paralog of CDK12 with a similar domain architecture and its kinase domain shares 90.5% identity to CDK12’s kinase domain^33^. Expression of the wild-type kinase domain of CDK13 also supported CCNK degradation. Mutating CDK13’s glycine 709 to glutamate or arginine (G709E or G709R), which is located at an equivalent position of glycine 731 in CDK12, also blocked CCNK degradation (Fig. S3D).

### HQ461 mediates interaction between CDK12 and DDB1

To characterize the mechanisms through which HQ461 triggers CCNK degradation, we used CRISPR/Cas9 to knock-in a sequence encoding an N-terminal 3xFLAG tag into *CDK12* and then immunoprecipitated 3xFLAG-CDK12 and its interacting proteins from cellular lysates treated with HQ461. Using western blotting, we found CDK12 stably associated with CCNK, doing so in an HQ461-independent manner. In contrast, DDB1 was only detected with the CDK12/CCNK complex isolated from lysates treated with HQ461 (Fig. 4A), suggesting that HQ461 mediates recruitment of CDK12/CCNK to DDB1. To recognize a substrate, DDB1-CUL4-RBX1 requires a type of substrate-specific receptor protein that known as a DDB1 CUL4 associated factor (DCAF). More than 60 human DCAFs have been discovered^34^. However, our genome-wide CRISPR-Cas9 screening of HQ461 resistance did not reveal any candidate DCAF (Fig. 1F), suggesting that HQ461-dependent degradation of CCNK may not require a DCAF. To test this, we next reconstituted CCNK polyubiquitination *in vitro* with purified recombinant ubiquitin, E1 (UBA1), E2 (UBE2G1 and UBE2D3), and E3 (CUL4-RBX1-DDB1) (Fig. S4A). CCNK in complex with wild-type CDK12 was efficiently polyubiquitinated in the presence of HQ461 (Fig. 4B). Omitting any enzyme from the reaction prevented CCNK polyubiquitination (Fig. S4B). CCNK in complex with CDK12 G731E or G731R was resistant to HQ461-induced polyubiquitination (Fig. 4B). Taken together, these results suggest that HQ461 promotes the recruitment of CDK12/CCNK to DDB1-CUL4-RBX1 for polyubiquitination.

**Figure 4.**
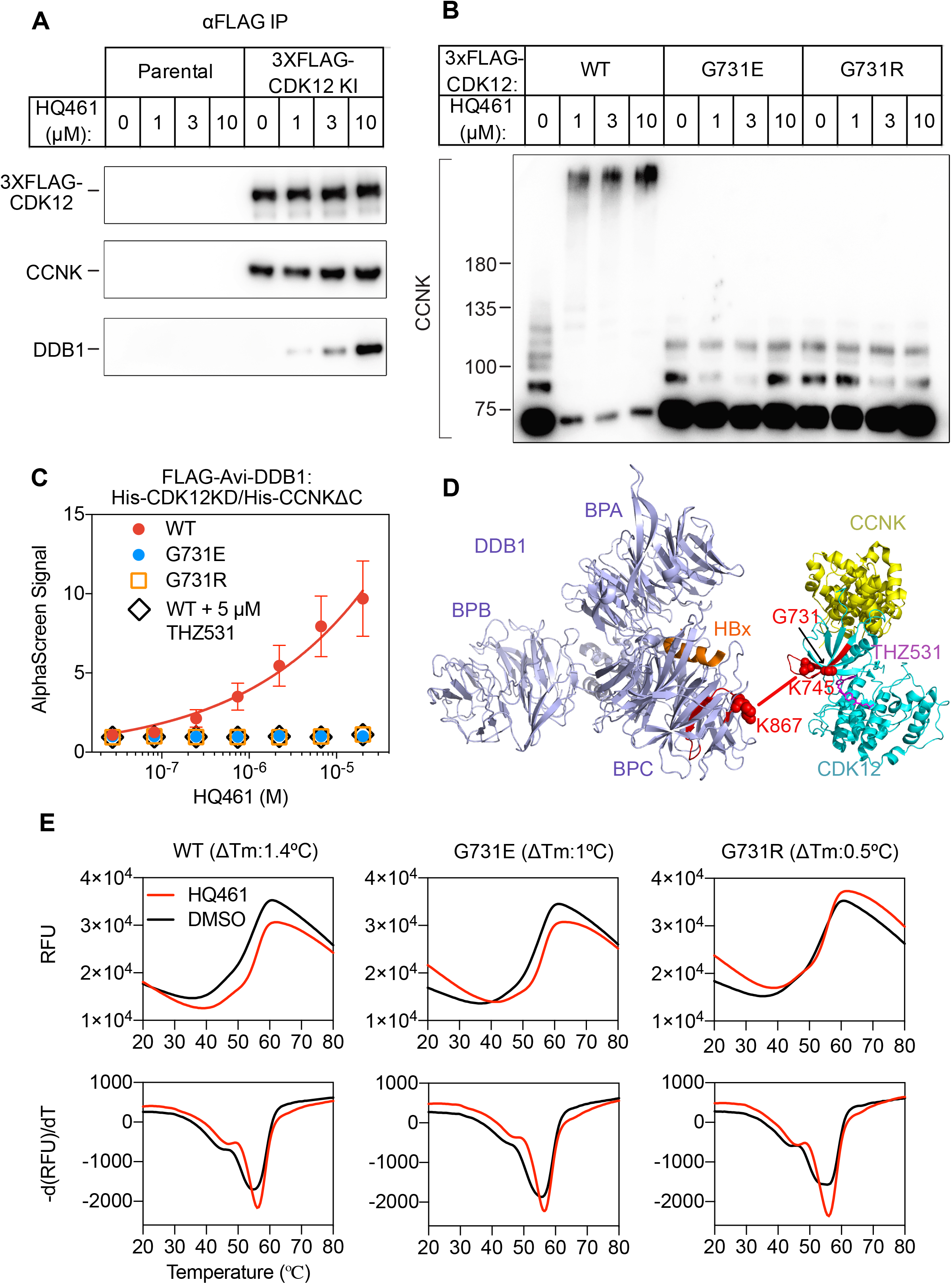
HQ461 functions as a molecular glue between CDK12 and DDB1. (A) Coimmunoprecipitation of DDB1 with 3xFLAG-CDK12/CCNK in the presence of HQ461. (B) In vitro ubiquitination of CCNK with recombinant ubiquitin, E1 (UBA1), E2 (UBE2G1 and UBE2D3), and E3 (DDB1-CUL4-RBX1). (C) Detection of HQ461-dependent interaction between FLAG-Avi-DDB1 and His-CDK12KD/His-CCNK∆C examined in an AlphaScreen assay. Error bars represent SEM from three technical replicates. (D) CXMS analysis identified one inter-protein cross-link between the DDB1(PDB code: 3i7h) lysine 867 and CDK12KD/CCNK∆C (PDB code: 5acb) lysine 745. (E) DSF analysis of CDK12KD/CCNK∆C (wild type and G731E/R mutants) in the presence of 50 μM HQ461 or vehicle DMSO.

### HQ461 functions as a molecular glue between CDK12 and DDB1

The apparent biochemical activity of HQ461 in mediating protein-protein interactions resembles previously reported molecular glues such as auxin, lenalidomide, and indisulam. To test if HQ461 also functions as a molecular glue, we purified FLAG-DDB1 as well as CDK12’s kinase domain (CDK12KD) in complex with the Cyclin box domain of Cyclin K (CCNK∆C). Using FLAG antibody-conjugated resin to pulldown FLAG-DDB1, we found that DDB1 formed a near-stoichiometric complex with CDK12KD/CCNK∆C, doing so in an HQ461-dependent manner (Fig. S4C). We further developed an AlphaScreen assay to measure the interaction between CDK12KD/CCNK∆C and DDB1, and observed that HQ461 induced the formation of a CDK12KD/CCNK∆C/DDB1 complex with an apparent EC_50_ of 1.9 μM. G731E and G731R mutants of CDK12KD/CCNK∆C did not form a complex with DDB1 even with 20 μM of HQ461 (Fig. 4C).

To map the HQ461-induced protein-protein interaction interface, we performed chemical cross-linking mass spectrometry (CXMS)^35^. Specifically, the HQ461-induced CDK12KD/CCNK∆C/DDB1 complex was cross-linked using two lysine-specific cross-linkers (DSS and BS3), followed by digestion with trypsin and LC-MS/MS analysis. Using pLink 2 software^36^, 41 pairs of cross-linked peptides were identified, out of which only one pair was derived from inter-protein cross-linking; the remainder were all derived from intra-protein cross-linking (Table S3). For this single inter-protein peptide pair, lysine 745 of CDK12 was cross-linked to lysine 867 of DDB1 (Fig. 4D).

### HQ461 binds to CDK12’s kinase domain

Like other kinases, CDK12’s kinase domain possesses an ATP-binding pocket sandwiched between an N-terminal β sheet-rich lobe and a C-terminal α helix-rich lobe^31,32^. Both lysine 745 and glycine 731 reside in the N-terminal lobe atop the ATP-binding pocket of CDK12, suggesting that HQ461 might bind to this pocket, creating a modified CDK12 protein surface to bind to DDB1. We tested this hypothesis by directly measuring HQ461 binding to CDK12KD/CCNK∆C using differential scanning fluorimetry (DSF) with a fluorescent dye Sypro Orange. We monitored the thermal denaturation of CDK12KD/CCNK∆C in the presence or absence of HQ461 and found that HQ461 stabilized CDK12KD/CCNK∆C by a 1.4 °C increase in its melting temperature (Tm) consistent with the model that HQ461 directly binds to CDK12KD/CCNK∆C (Fig. 4E). We next evaluated whether HQ461’s binding to CDK12KD/CCNK∆C was affected by G731E/R mutations. By DSF, HQ461 caused 1 °C and 0.5 °C shifts in the Tm of G731E and G73R mutants of CDK12KD/CCNK∆C, respectively. Although the magnitude of HQ461-dependent Tm shifts is smaller for G731E/R relative to wild type protein, these results suggests that HQ461 is still capable of binding to G731E and G731R mutant CDK12 kinase domain.

We next used DSF to test if HQ461 also binds to DDB1. DDB1 is composed of three β propeller domains (BPA, BPB, and BPC) (Fig. 4D)^37^. The residue lysine 867 on DDB1 that is cross-linked to CDK12 in the CXMS experiment is located in the BPC domain of DDB1, which is known to interact with a promiscuous α-helical motif present in DCAF proteins and viral proteins such as the hepatitis B virus X protein (HBx)^38^. We found that a peptide corresponding to the α-helical motif of HBx increased the thermal stability of recombinant DDB1 without its BPB domain (DDB1∆BPB) by a 2 °C increase in Tm (Fig. S5A). As a negative control, HBx peptide carrying an arginine to glutamate substitution, which is known to abrogate DDB1 binding^38^, did not cause a change in the Tm of DDB1∆BPB. Similar with the negative control, HQ461 did not change the thermal stability of DDB1∆BPB (Fig. S5A). These results suggest that HQ461 does not directly bind to DDB1.

THZ531 (Fig. S5B) is a specific covalent inhibitor of CDK12/CDK13 by irreversibly targeting a cysteine uniquely present in CDK12/13’s kinase domains^39^. Using the AlphaScreen assay, we found that THZ531 inhibited HQ461-dependent formation of the CDK12KD/CCNK∆C/DDB1 complex (Fig. 4C). Dinaciclib (Fig. S5B) is a non-covalent CDK inhibitor targeting CDK1, 2, 5, 9, 12, 13^40^. Neither THZ531 nor dinaciclib treatment caused CCNK degradation (Fig. S5C). Furthermore, treating cells with THZ531 or dinaciclib prior to HQ461 treatment prevented CCNK degradation, clearly suggesting that THZ531, dinaciclib, and HQ461 bind to the same ATP-binding pocket in CDK12’s kinase domain. These results confirm HQ461’s engagement with CDK12’s kinase domain *in vivo*, and highlight HQ461’s unique ability to induce CCNK degradation.

### Structure-activity relationship study of HQ461

To dissect the chemical basis for optimizing HQ461’s molecular-glue activity, we conducted a thorough structure-activity relationship study of HQ461 (Fig. 5 and S6). We made a CCNK-luc reporter that converts the degradation of CCNK into a reduction in luciferase activity (Fig. S7). We then measured the ability of each synthesized analog in reducing CCNK-luc activity at 10 μM relative to DMSO control (maximal degradation, Dmax). For analogs with a Dmax greater than 50%, we further performed dose response analysis to measure their half-maximal CCNK-degradation concentrations (DC_50_) (Fig. 5 and S7).

**Figure 5.**
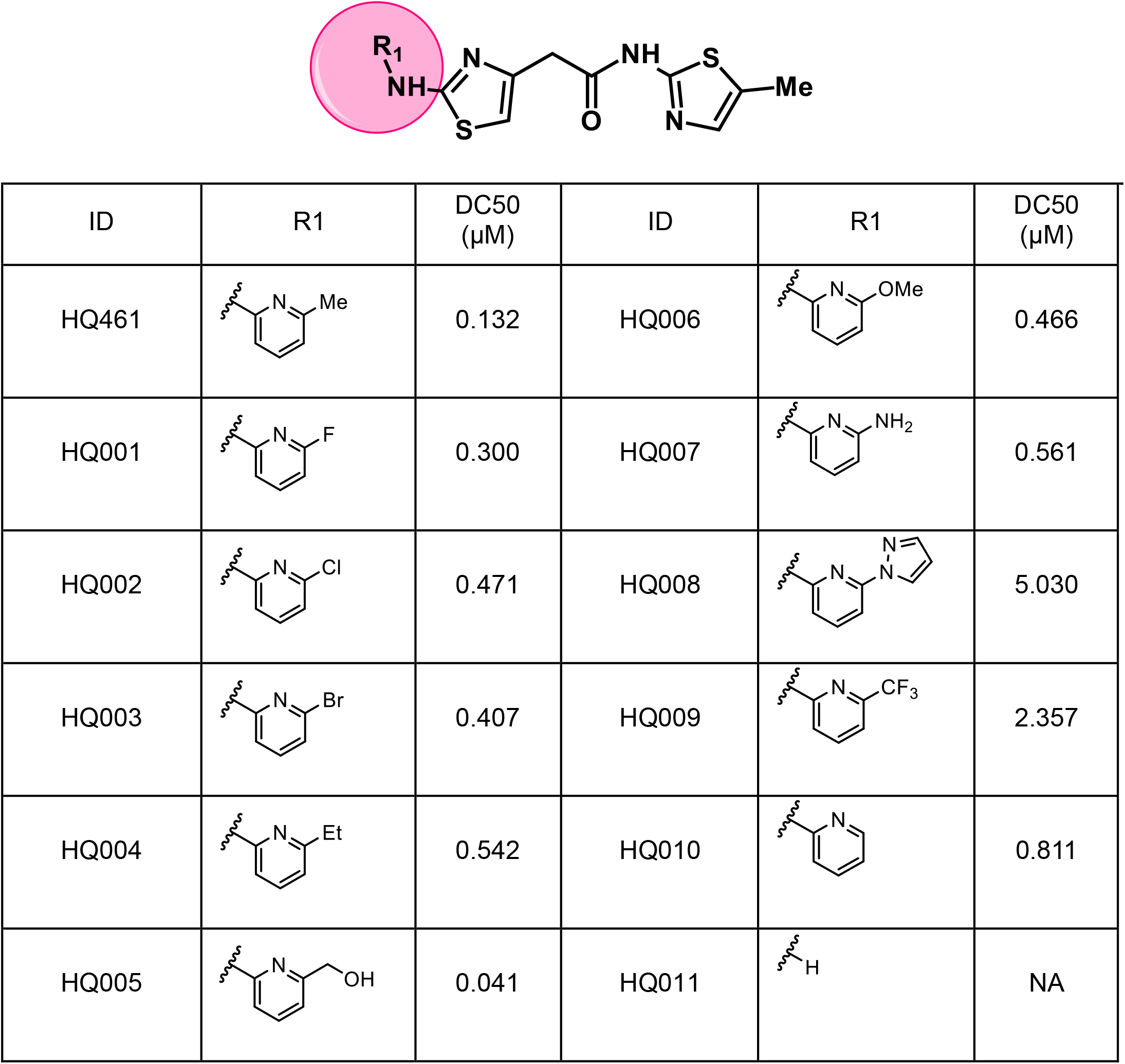
Structure-activity relation of HQ461 analogs. Half maximal CCNK-degradations are reported for each analog. CCNK degradation was measured using a CCNK-luc reporter.

We first modified the aminopyridinylthiazol scaffold either by changing pyridine to other substituted heterocycles or by elongating the linker to increase the overall size of the molecule (Fig. S6). Specifically, we changed the pyridine to small-size group hydrogen (HQ011) and acetyl (HQ019), and replaced the linker between pyridine and aminothiazol to amide (HQ014). These modifications resulted in losses in activity. We also made changes to the 5-methylthiazol-2-amine moiety of HQ461 by incorporating several substitutions (HQ015, 016, and 017) and discovered that this moiety was crucial for activity. We therefore maintained this pharmacophore and incorporated a variety of substitutions on the pyridine ring (Fig. 5). The elimination of the methyl group (HQ010) on pyridine ring resulted in a 6-fold higher DC_50_ value, suggesting an important steric effect of the 5-position on the pyridine ring. Consistent with this hypothesis, the incorporation of a bulky ring structures on pyridine’s 5-position (HQ008) led to a 38-fold increase in DC_50_. In addition, substitution of the 5-position with F, Cl, Br, ethyl, methoxyl, and trifluoromethyl (HQ001-004, 006, and 009) all diminished the potency.

Finally, a beneficial effect of polarity is evidenced by a 3-fold increase of potency in HQ005 (DC_50_: 0.041 μM) in which the methyl group in HQ461 was replaced by a hydroxymethyl. We speculate that this hydroxyl group may interact favorably with polar residues in the pocket of CDK12 kinase domain. In summary, our SAR efforts resulted in an improved compound HQ005 with a hydroxymethyl moiety at the 5-position of the pyridine ring and displayed a DC_50_ of 41 nM. Further SAR optimization and lead development including better pharmacokinetic and pharmacodynamic properties will be a focus of future studies.

### HQ461-dependent degradation of CCNK causes decreased NRF transcription and cell death

In addition to cytotoxicity, HQ461 also affects the transcription of NRF2 (Fig.S1C). We next examined if these two molecular phenotypes were consequences of CCNK depletion. CCNK forms two distinct complexes with CDK12 and with CDK13^27,28^, and is essential for the kinase activity of both complexes. Inactivation of CCNK therefore blocks the functions of both CDK12 and CDK13. Indeed, treating cells with the dual CDK12/CDK13 inhibitor THZ531 also caused a reduction of the NRF2 level (Fig. S8A).

The C-terminal domain of RNA polymerase II’s largest subunit is composed of 52 repeats of with the consensus heptad sequence YSPTSPS. Serine 2 of heptad repeat is a known phosphosubstrate of CDK12/13^27^. Both THZ531 and HQ461 reduced Serine 2 phosphorylation of POLII CTD without affecting total POLII levels (Fig. 6A and S8B). Inactivation of CDK12 is known to preferentially affect the expression of genes involved in DNA damage response (DDR)^27,40^. We used RT-qPCR to monitor the mRNA levels of *BRCA1*, *BRCA2*, *ATR*, and *ERCC4*, four DDR genes known to be downregulated by CDK12 inhibition. HQ461 treatment resulted in a time-dependent reduction in the mRNA levels of all four tested genes (Fig. 6B). Finally, we tested if genetic depletion of CCNK causes cell death. Using a colony formation assay, an sgRNA targeting *CCNK* in A549 cells resulted in reduced cell viability. The inhibitory effect of *CCNK*-targeting sgRNA could be reversed by expressing a cDNA sequence containing an sgRNA-resistant variant of *CCNK*, confirming that the observed inhibitory effect results from CCNK inactivation (Fig. 6C). Taken together, these results support the conclusion that NRF2 downregulation and cytotoxicity all result from HQ461-mediated depletion of CCNK (Fig. 6D).

**Figure 6.**
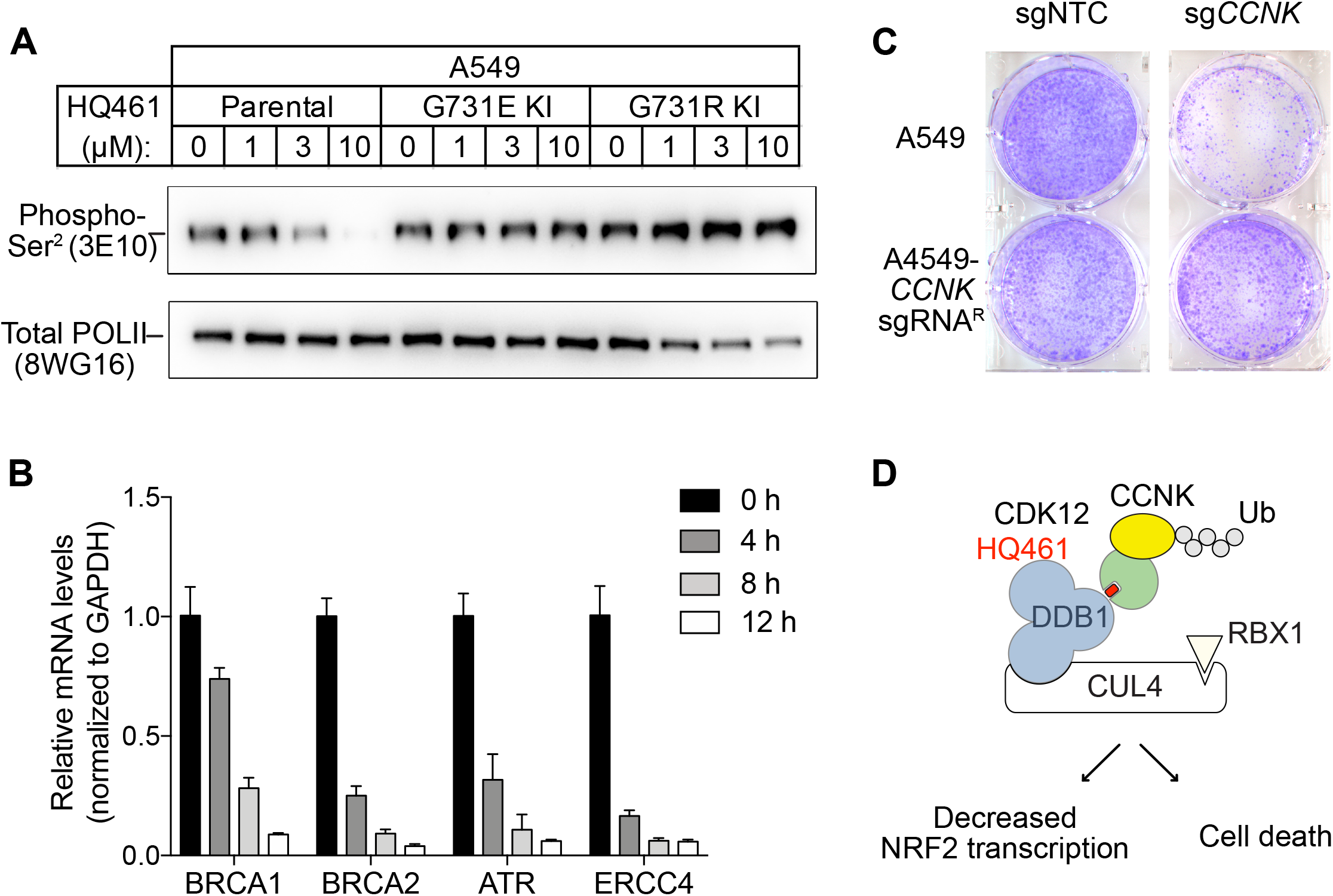
HQ461-mediated CCNK degradation causes reduced NRF2 transcription and cell death. (A) Western blotting to examine levels of POLII CTD serine 2 phosphorylation and total POLII following HQ461 treatment. (B) RT-qPCR to examine levels of BRCA1, BRCA2, ATR, ERCC4 mRNAs following 10 μM HQ461 treatment. Error bars represent standard deviations (SD) from three technical replicates. (C) Depletion of CCNK by CRISPR reduces colony formation in parental A549 cells but not in A549 cells expressing an sgRNA-resistant *CCNK* cDNA. (D) A model of HQ461’s mechanism of action.

## Discussion

HQ461 was discovered in high-throughput screening of small molecules that inhibited NRF2 activity (Fig. 1A and 1C). NRF2 is a basic leucine zipper transcription factor that regulates the expression of genes encoding xenobiotic-detoxification or antioxidant enzymes^15^. In the absence of oxidative or electrophilic stress, NRF2 is degraded by the ubiquitin-proteasome system through binding to the E3 ubiquitin ligase KEAP1-CUL3-RBX1^15^. In lung and liver cancer patients, *KEAP1* loss-of-function or *NRF2* gain-of-function mutations are frequently observed^16,17^, suggesting that over-active NRF2 is a potential therapeutic target for these cancers. To our surprise, when we used CRISPR-Cas9 to inactivate *NRF2* in A549 cells, we did not observe an overt growth defect; an isolated *NRF2* knockout A549 clone proliferated at a comparable rate relative to parental A549 cells. These results suggest that NRF2 activity is not essential for optimal growth of A549 cells under optimal tissue-culture conditions. Whether NRF2 activity is required for the *in vitro* proliferation of other *KEAP1* mutant cancer cell lines will be investigated in the future.

Based on our biochemical and biophysical data, we propose a model in which HQ461 binds to CDK12’s ATP-binding pocket to generate an altered surface on CDK12 to interact with the BPC domain of DDB1. This model is supported by multiple lines of evidence. First, DSF experiments revealed direct binding of HQ461 to the CDK12/CCNK complex instead of DDB1 (Fig. 4F and S5A). Second, known inhibitors of CDK12 that bind to CDK12’s ATP-binding pocket impeded HQ461’s activity in inducing CDK12-DDB1 interaction and CCNK degradation (Fig. 4C and S5D). Third, chemical cross-linking mass spectrometry analysis mapped the HQ461-induced protein-protein interaction interface to be between CDK12’s kinase domain and DDB1’s BPC domain (Fig. 4D). Interestingly, the hot spot glycine 731 for HQ461-resistant mutations is located in the vicinity of this interface. Replacing glycine (a neutral residue without a side chain) with glutamate or arginine (charged residues with bulky side chains) may cause steric hindrance and/or disruption of hydrophobic stacking to destabilize CDK12 and DDB1’s interaction interface. High-resolution structural analyses of a CDK12-DDB1-HQ461 complex is needed to completely resolve the basis for HQ461’s molecular-glue activity.

Previously known molecular-glue degraders all interact with a substrate-specific receptor of Cullin RING ligases (CRL) to exert their functions. Auxin, thalidomide, and indisulam bind to TIR1, cereblon, and DCAF15, respectively^1,5,8^. HQ461 engages the CRL in an unprecedented manner by directly interacting with the adaptor protein DDB1. CRLs catalyze polyubiquitination of their protein substrates by positioning them in a reactive zone. CDK12 appears to be outside this reactive zone, because HQ461 did not cause CDK12 ubiquitination (Fig. 3D). In contrast, CDK12’s partner protein CCNK seems to be optimally positioned for polyubiquitination. These findings suggest that HQ461-bound CDK12 is functionally equivalent to a substrate-specific receptor protein for CRL. Interestingly, several viruses have evolved a similar strategy to use viral proteins as facultative substrate-specific receptors for DDB1 to degrade host restriction factors^37,38^.

CDK12 and CDK13 are two structurally similar kinases that has the ability to phosphorylate serine 2 of the POLII CTD heptad repeats^31–33^. To execute their functions, CDK12 and CDK13 form two separate complexes with CCNK^27,28^. Consistent with this notion, HQ461-mediated depletion of Cyclin K phenotypically mimics dual inhibition of CDK12 and CDK13 by CDK12/13-selective inhibitor THZ531 (Fig. S8A). CDK12 inactivation is known to cause reduced POLII elongation and premature polyadenylation of select genes enriched in the DNA damage response pathway, resulting in a BRCAness phenotype^27,40,41^. Several studies have shown that inhibiting CDK12 can sensitize cancer cells to PARP inhibitors^40,42,43^. It will be interesting to test whether HQ461, by depleting CCNK, also enhance the therapeutic efficacy of PARP inhibitors for cancer treatment.

## Acknowledgements

We thank D. Nijhawan, L. Sun, S. Zheng, F. Lu for cell lines and reagents, and J. Snyder for critically reading and editing the manuscript. This work was supported by institutional grants from the Chinese Ministry of Science and Technology and Beijing Municipal Commission of Science and Technology. The funders had no role in study design, data collection and interpretation, or the decision to submit the work for publication.

---

**Figure S1.**
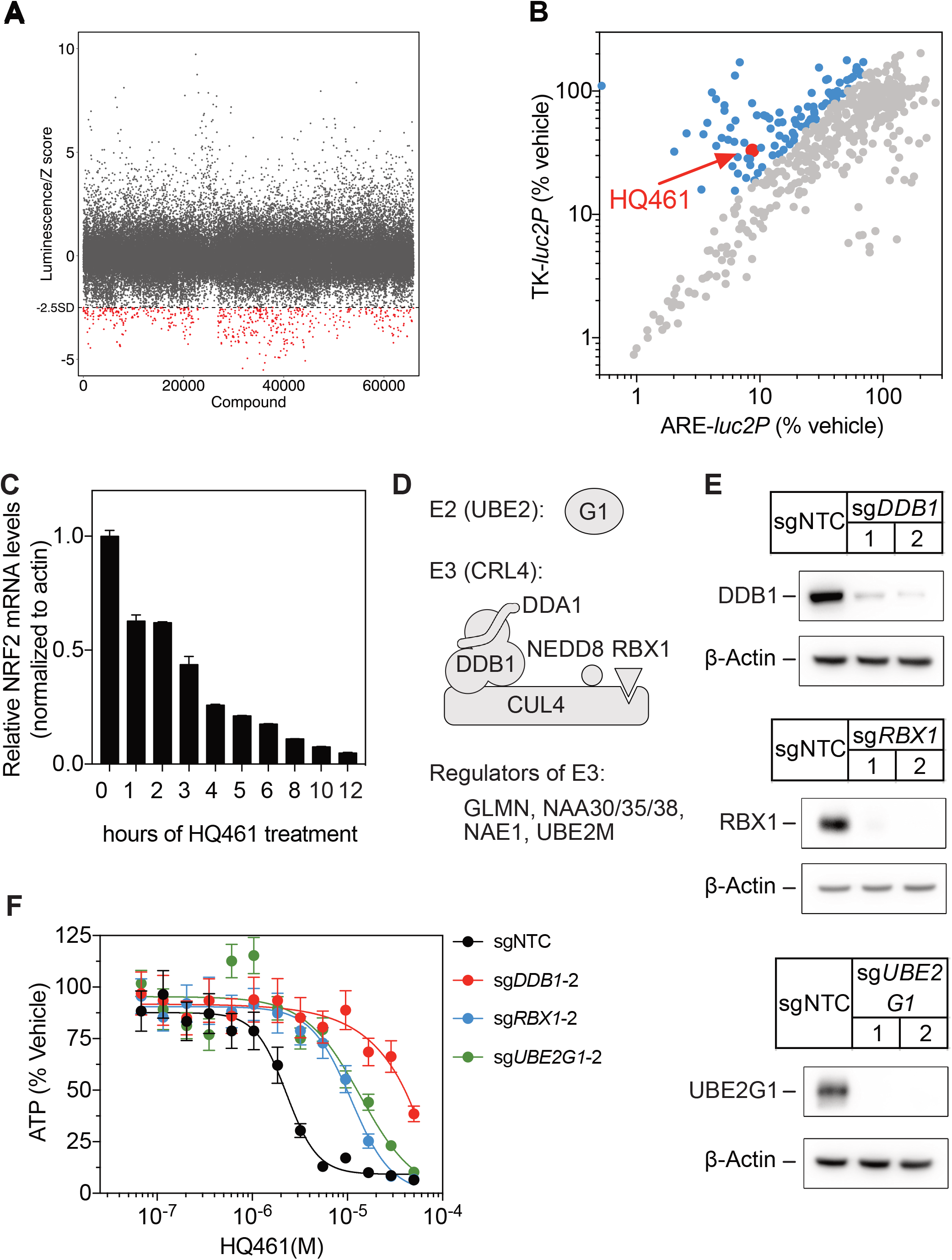
High-throughput chemical screening and CRISPR-Cas9 genetic screening demonstrate that HQ461 requires DDB1-CUL4-RBX1 for its cytotoxicity. (A) Primary screen results. The ARE-luc2p reporter cell line was treated for 12 hours with library compounds at 10 μM. Each screening plate was individually Z-score normalized, and all the Z-scores were aggregated. The dotted line represents the cutoff (Z-score < −2.5). Red dots mark hits from the primary screen. (B) Counter screen results. Hits from the primary screen were tested in both ARE-*luc2P* and TK-*luc2P* reporter cell lines at 10 μM for 12 hours. Blue dots represent compounds selectivity inhibiting ARE-*luc2P* relative to TK-*luc2P*. (C) Analysis of *NRF2* mRNA level in A549 cells from a time course of HQ461 (10 μM) treatment. Error bars represent SD from three technical replicates. (D) Classification of top-ranking genes into functional categories in the ubiquitin protein system. (E) Verification of successful depletion of DDB1, RBX1, and UBE2G1 by individual sgRNAs. (F) Measurement of the HQ461 IC_50_ on the viability of A549 cells expressing non-targeting control (NTC) or a second set of sgRNAs targeting *DDB1*, *RBX1*, or *UBE2G1* that are independent of the sgRNAs used for the experiments presented in Figure 1G. Error bars represent SEM from three biological replicates.

**Figure S2.**
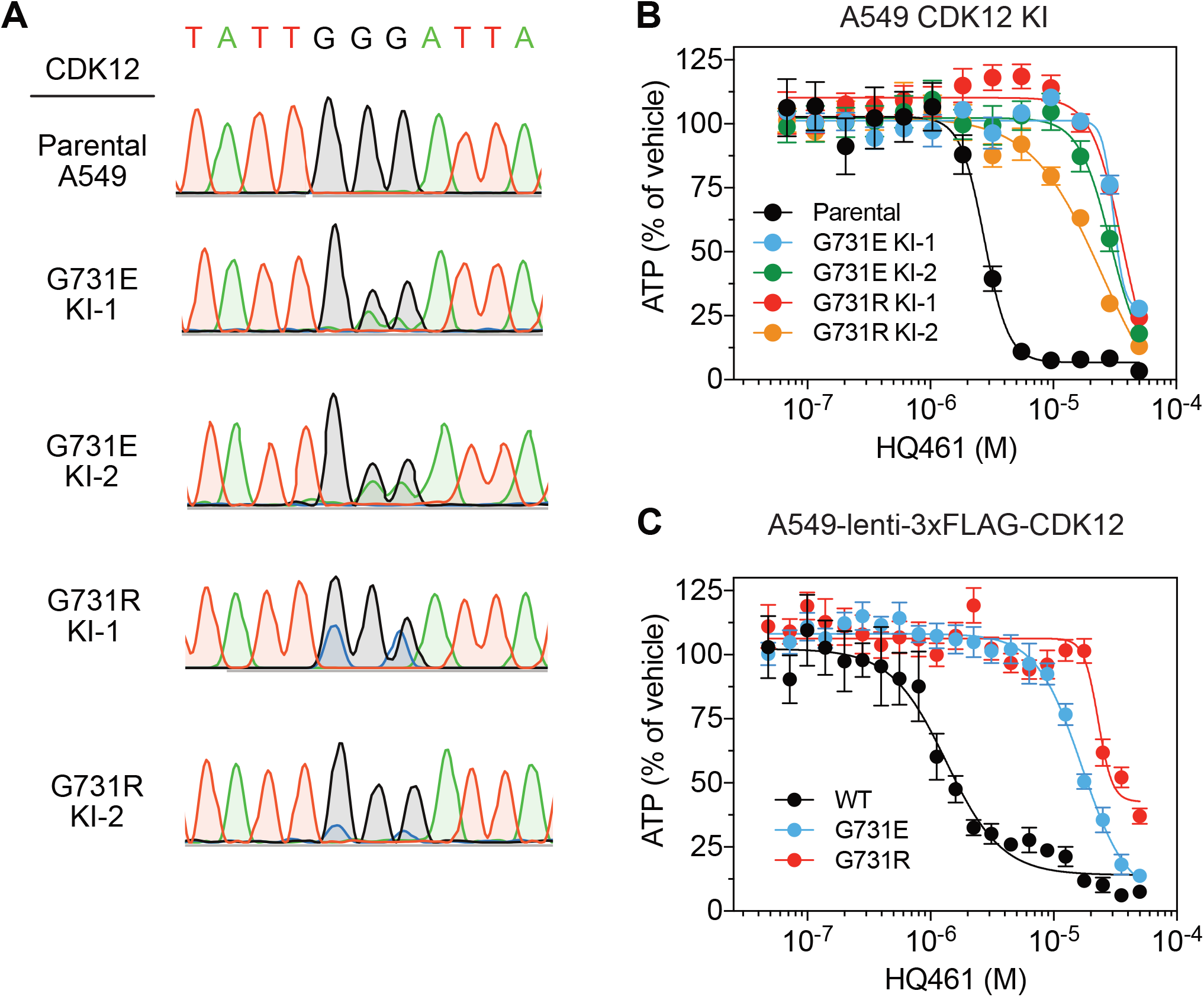
The *CDK12* G731E and G731R mutations confer HQ461 resistance. (A) Sanger sequencing verification of GGG to GAA and GGG to CGC mutations introduced by CRISPR-Cas9 mediated homology dependent repair. (B) Measurement of the HQ461 IC_50_ on the viability of CDK12 G731E or G731R knock-in A549 cells. Error bars represent SEM from three biological replicates. (C) Measurement of the HQ461 IC_50_ on the viability of A549 cells stably expressing wild-type or G731E/R mutant CDK12. Error bars represent SEM from three biological replicates.

**Figure S3.**
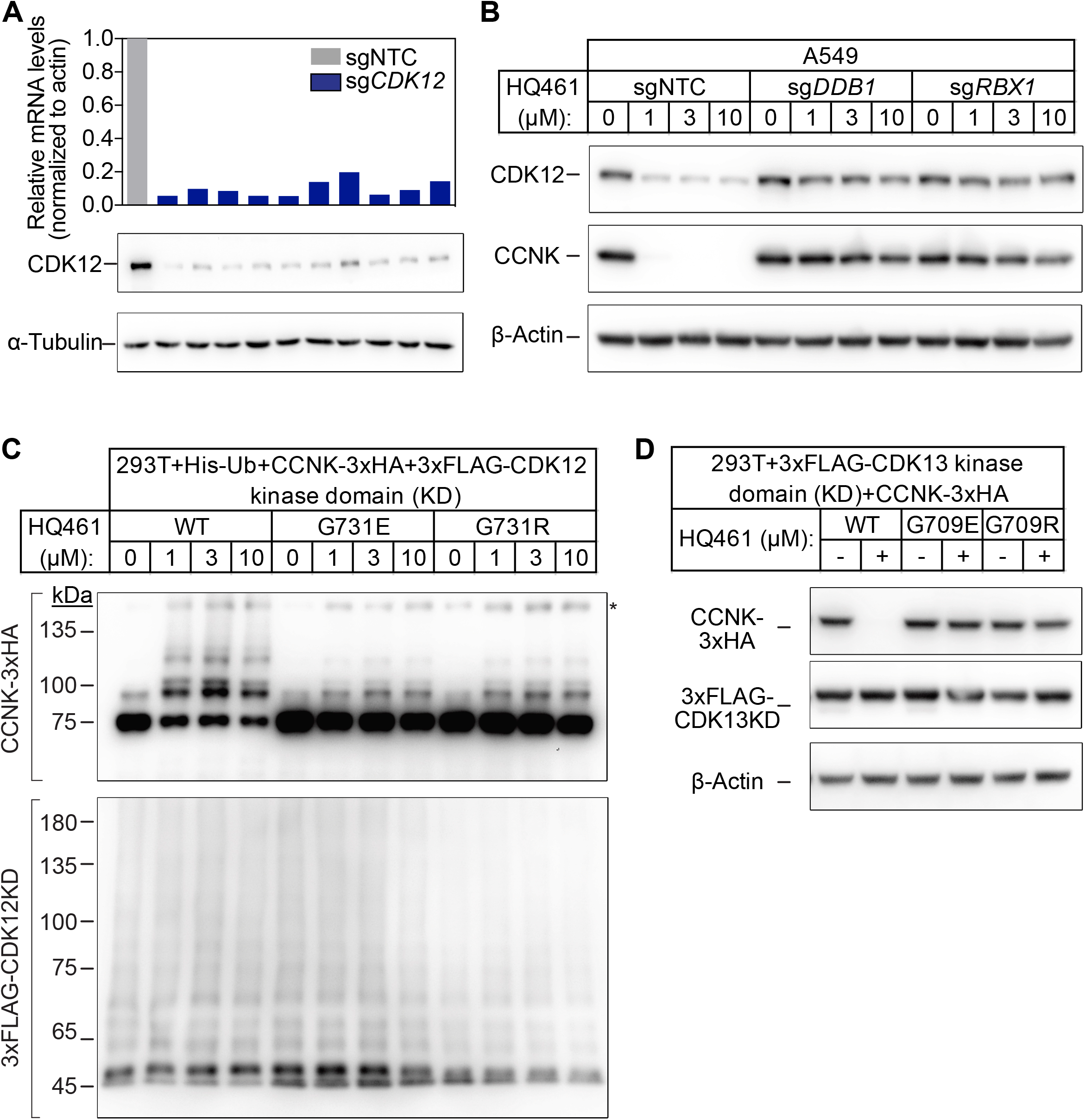
Both CDK12 and CDK13 support HQ461-induced CCNK degradation. (A) Measurement of CDK12 (mRNA and protein levels) following CDK12 CRISPRi. (B) Effect of DDB1 and RBX1 depletion on CCNK and CDK12 protein levels in A549 cells treated with HQ461. (C) HQ461 triggers in vivo polyubiquitination of CCNK in complex with the wild-type CDK12 kinase domain but not with G731E or G731R mutant kinase domains. (D) Wild-type CDK13 kinase domain but not G709E or G709R mutant kinase domain is sufficient for HQ461-mediated destabilization of CCNK.

**Figure S4.**
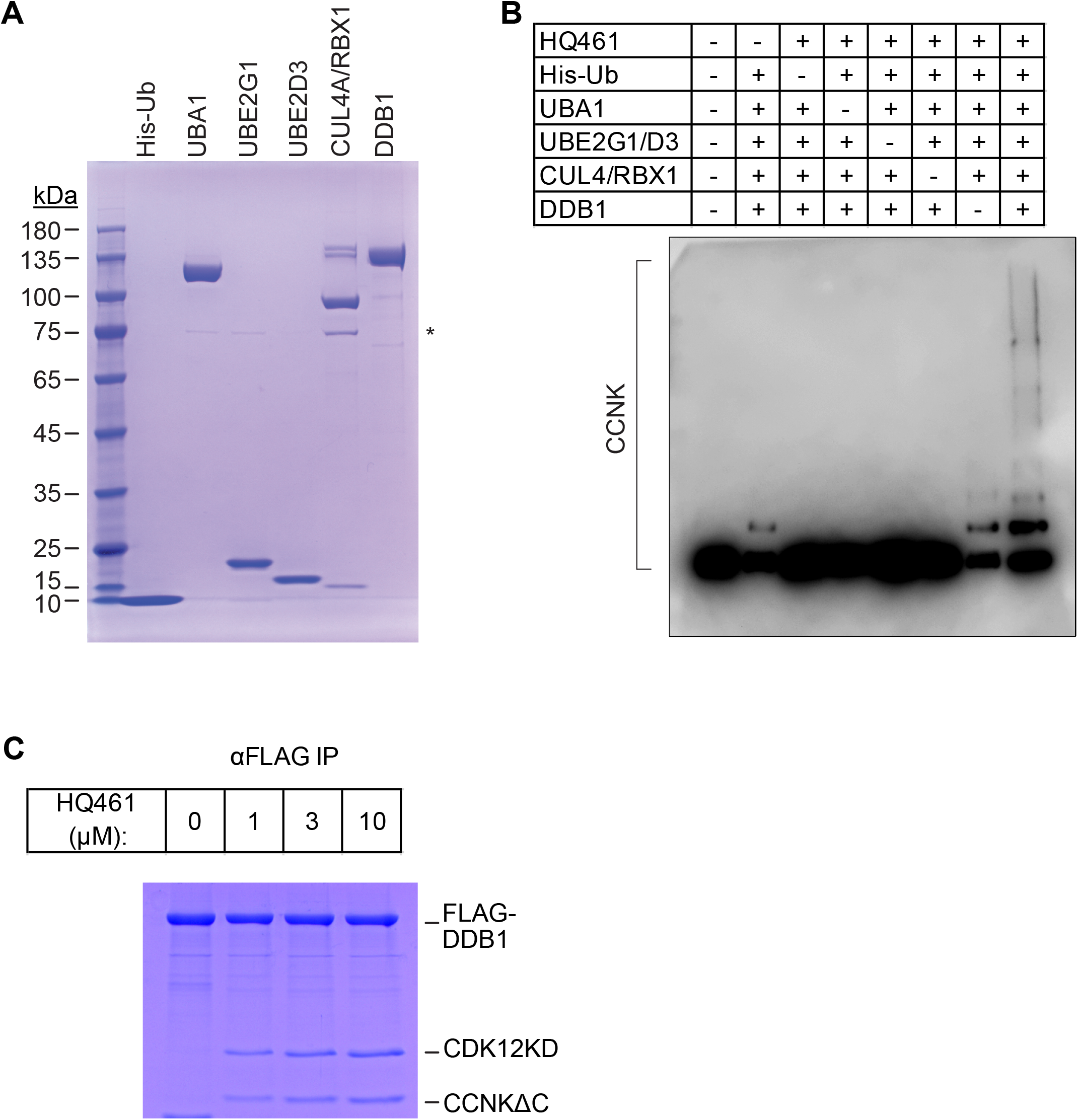
Biochemical characterization of HQ461’s molecular glue activity. (A) Coomassie blue staining of recombinant proteins used for an in vitro ubiquitination assay. The asterisk indicates contaminating HSP70. (B) In vitro ubiquitination of CCNK with recombinant ubiquitin (His-Ub), E1 (UBA1), E2 (UBE2G1 and UBE2D3), and E3 (DDB1-CUL4-RBX1). (C) HQ461-dependent formation of a FLAG-DDB1/CDK12KD/CCNK∆C complex.

**Figure S5.**
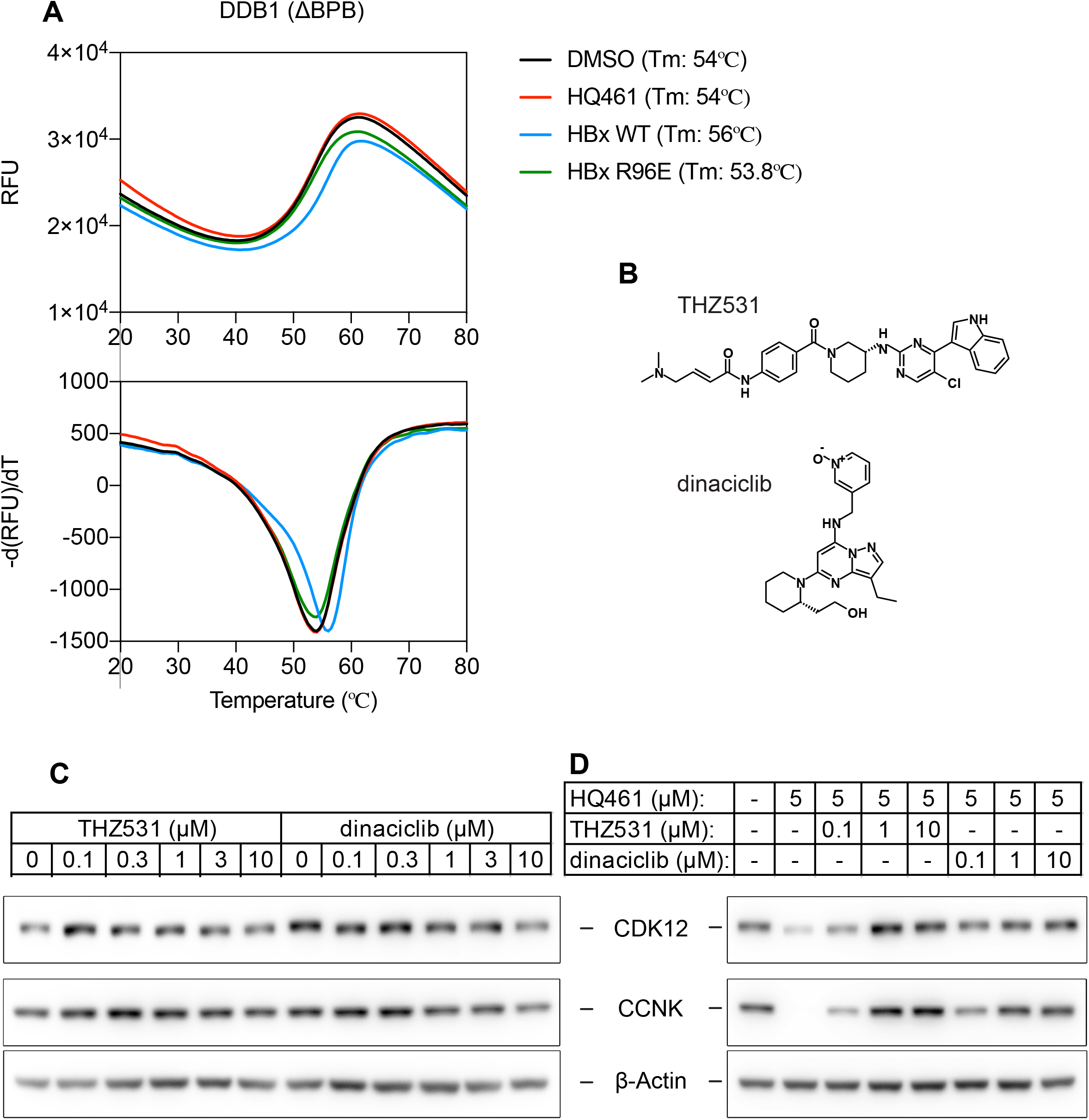
HQ461 binds to the kinase pocket of CDK12. (A) DSF analysis of DDB1∆BPB in the presence of DMSO, 50 μM HQ461 or HBx peptide (WT or R96E mutant). (B) Chemical structures of THZ531 and dinaciclib. (C) Effect of THZ531 and dinaciclib on CCNK and CDK12 protein levels in A549 cells. (D) Effect of THZ531 and dinaciclib on the CCNK and CDK12 protein levels in A549 cells treated with HQ461.

**Figure S6.**
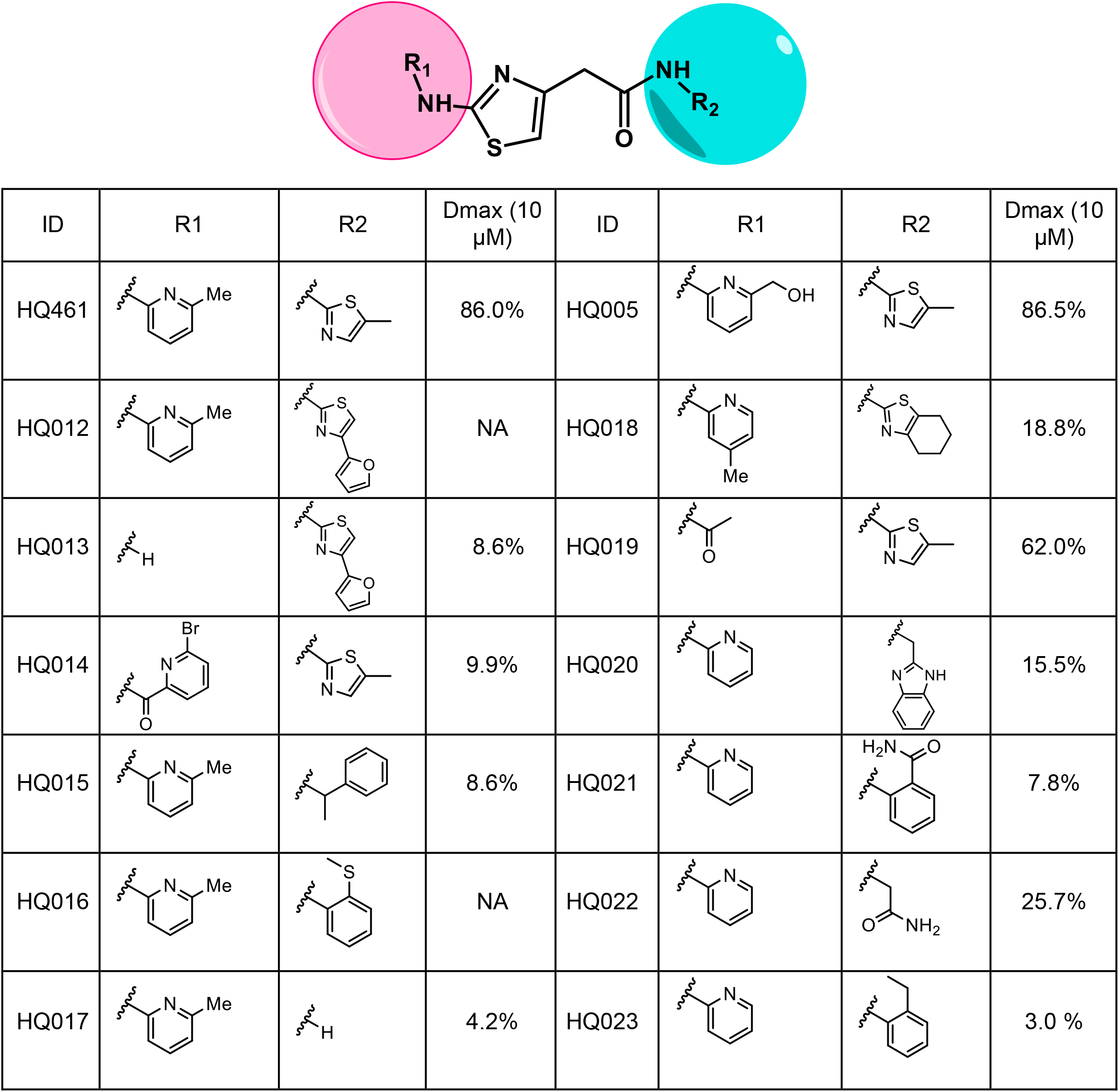
Structure-activity relation of HQ461 analogs. Dmax represents the percentage of CCNK degradation with 10 μM of test compound treatment relative to DMSO control. CCNK degradation was measured using a CCNK-luc reporter.

**Figure S7.**
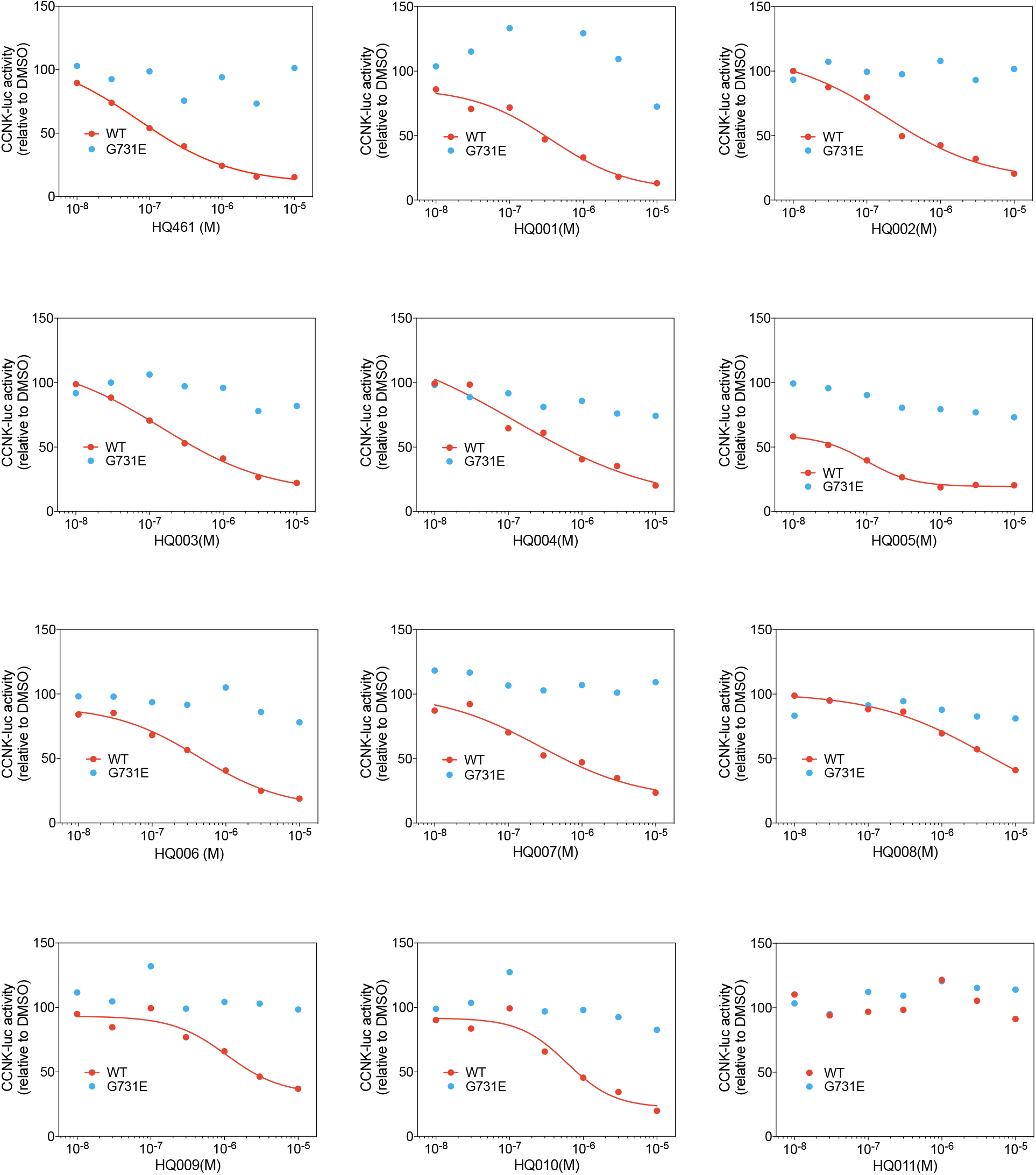
Dose response curves of HQ461 analogs in reducing CCNK-luc activity. The CCNK-luc reporter co-transfected with CDK12KD with the G731E mutation verifies that CCNK-luc degradation is on-target.

**Figure S8.**
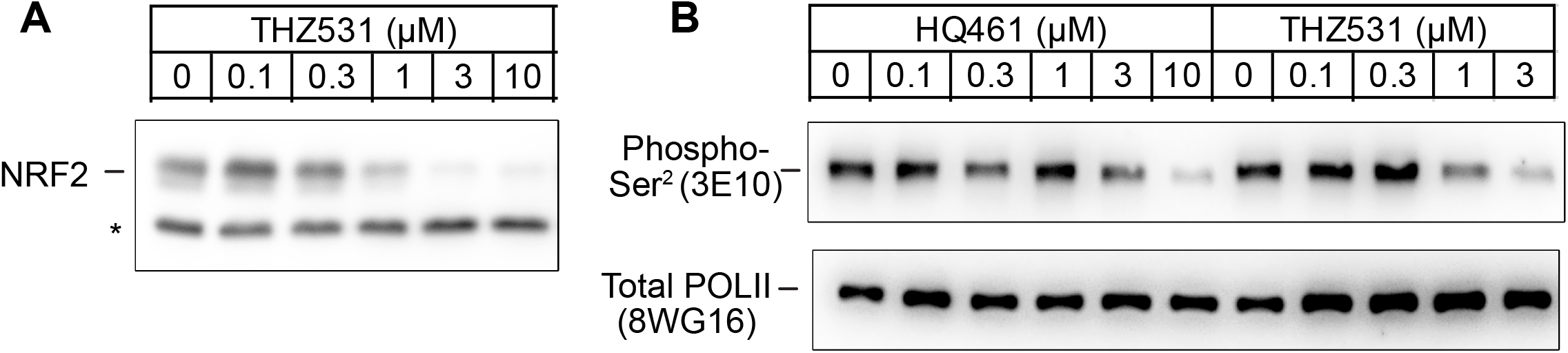
Both HQ461 and THZ531 reduce NRF2 protein levels and POLII CTD serine 2 phosphorylation. (A) Western blotting of NRF2 following THZ531 treatment for 8 hours. Loading control β-Actin is as in Figure S5C. (B) Western blotting of POLII CTD serine 2 phosphorylation and total POLII following HQ461 or THZ531 treatment for 8 hours.

## Materials and methods

### Cell culture

The human cell lines A549, HCT-116, and HEK293T were obtained from Dr. Deepak Nijhawan’s lab at University of Texas Southwestern Medical Center. Free style 293-F was a gift from Dr. Linfeng Sun at China University of Science and Technology. The insect cell lines Sf9 and High Five were gifts from Dr. Sanduo Zheng from National Institute of Biological Sciences, Beijing. The identities for A549 and HCT-116 were confirmed by short tandem repeat (STR) analysis. All cell lines were confirmed to be mycoplasma free on a weekly basis using a PCR-based assay with primers: 5’-GGGAGCAAACAGGATTAGA TACCCT-3’ and 5’-TGCACCATCTGTCACTCTGTTAACCTC-3’. Regular adherent cell culture methods were used to culture A549, HCT-116, and HEK293T cells in tissue-culture incubators with 5% CO_2_ at 37 °C. A549 were grown in RPMI-1640 medium with 10% fetal bovine serum (FBS) and 2 mM L-glutamine. HCT-116 and HEK293T cells were grown in DMEM medium with 10% FBS and 2 mM L-glutamine. Regular suspension cell culture methods were used to grow 293-F, Sf9, and high five cells. Free style 293-F cells were grown in SMM 293-TII expression medium (Sino Biological, M293TII) in a shaker incubator at 150 rpm, 37 °C, 5% CO_2_. Sf9 and High Five cells were cultured in ESF 921 medium (Expression Systems, 96-001-01) in a shaker at 140 rpm, 27 °C.

### Chemicals

Bortezomib (CAS No. 179324-69-7), THZ531 (CAS No. 1702809-17-3), and dinaciclib (CAS No. 779353-01-4) were purchased from Targetmol. MLN4924 (CAS No. 905579-51-3) was purchased from Selleckchem. All of these chemicals were prepared as 10 mM stocks in DMSO (CAS: 67-68-5) purchased from Sigma and further diluted in DMSO to the desirable concentrations. HBx peptides (WT: ILPKVLHKRTLGLS; R96E: ILPKVLHKETLGLS) were synthesized by ChinaPetides with a purity of 96%.

### Construction of ARE-*luc2P* and TK-*luc2P* reporter cell lines

A549 cells were transiently transfected with the plasmid pGL4.37[luc2P/ARE/Hygro] (Promega) and selected with 500 μg/ml of hygromycin to obtain a stable cell line harboring the ARE-*luc2P* reporter. TK-*luc2P* was cloned into the pLVX-IRE-Puro backbone by replacing its CMV promoter and multiple cloning site with HSV-TK promoter fused to *luc2P*. The resulting pLVX-TK-*luc2P*-IRES-Puro plasmid was packaged into lentivirus to transduce A549 cells. Stable cell lines were obtained by selection of transduced cells with 2 μg/ml of puromycin.

### High-throughput small-molecule screening

A screening library with 65,790 small molecules (Life Chemicals) were used in primary screening. The screening procedures were as follows. Seven hundred A549 ARE-l*uc2P* cells in 50 μL of medium were plated per well in 384-well flat clear bottom white polystyrene TC-treated microplates (Corning) and allowed to attach to plates overnight. Compounds from the screening library were added to the cells with Biomek FXP automated workstation at a final concentration of 10 μM. Twelve hours later, ARE-*luc2P* activities were measured with Bright-Glo luciferase assay system (Promega) following vendor instructions. Luminescence was measured on a EnVision multimode plate reader (PerkinElmer). Luminescence data on each plate was Z-score normalized using the equation 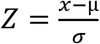, where x is the luminescence value of a well, μ is the mean and σ is the standard deviation of all values on one plate. Z scores from all the plates were aggregated and a cutoff (Z-score < −2.5) was used to select 515 hits from the primary screening. These hits were picked from the screening library and used for counter screening using both A549 ARE-*luc2P* and A549 TK-*luc2P* cells.

Normalization to negative control (DMSO) was used to calculate a % of inhibition value for every compound on the two *luc2P* reporters. A ratio (% inhibition of TK-luc2P / % inhibition of ARE-luc2P) greater than 2.5 was used to identify compounds with ARE-*luc2P* selectivity.

### Cell viability assay

Seven hundred A549 cells or 1,000 HCT-116 cells in 50 μL of medium were plated per well in 384-well flat clear bottom white polystyrene TC-treated microplates (Corning). After overnight attachment, cells were dosed with a serial dilution of HQ461 with a D300e digital dispenser (Tecan). Cell survival was measured 72 hours later using CellTiter-Glo luminescent cell viability assay kit (Promega) following vendor instructions. Luminescence was recorded by EnVison multimode plate reader (PerkinElmer). IC50 was determined with GraphPad Prism using baseline correction (by normalizing to DMSO control), the asymmetric (five parameter) equation, and least squares fit.

### Generation of A549*NRF2* knockout cells

An sgRNA sequence targeting *NRF2* (5’-TATTTGACTTCAGTCAGCGA-3’) was cloned into the lentiCRISPR v2 vector (Addgene #52961). Lentiviral packaging was performed by co-transfecting the resulting plasmid with psPAX2 (Addgene 12260) and pMD2.G (Addgene 12259) into 293T cells. Medium collected from transfected 293T cells was used to infect A549 cells followed by selection with 5 μg/ml of puromycin. Cells surviving puromycin selection were then plated sparsely on a 10-cm plate and clones were isolated. *NRF2* knockout clones were identified by NRF2 western blotting.

### Pooled genome-wide CRISPR-Cas9 sgRNA screening of HQ461 resistance

The human CRISPR knockout pooled library (Brunello)^18^ was a gift from Dr. Feiran Lu at University of Texas Southwestern Medical Center. The sgRNA library was packaged into lentiviral vector for delivery into A549 cells as described^44^. The screening parameters were as follows. Thirty million A549 cells were infected at a multiplicity of infection (MOI) of ~0.3. Infected cells were passaged every two days in the presence of 10 μg/ml of puromycin for one week with a population size of at least 200 million. Afterwards, A549 cells transduced with the sgRNA library were passaged every two days in the presence of 4 μM HQ461 or the vehicle DMSO (0.1% v/v) for three weeks with a population size of at least 10 million. Two biological replicates were performed for both HQ461 and DMSO treatment.

### Isolation of genomic DNA

Cells were lysed in 10 mM Tris-HCl, pH 8.0, 100 mM EDTA, 0.5% SDS, 200 μg/ml RNase A and incubated at 37 °C for 1 hour followed by the addition of 1mg/ml of proteinase K for an overnight incubation at 50 °C. The resulting lysates were extracted three times with phenol solution equilibrated with 10 mM Tris-HCl, pH 8.0, 1 mM EDTA (Sigma P4557) mixed with chloroform (1:1 v/v) using tubes with phase lock gel (Tiangen WM5-2302831). The aqueous phase was then mixed with 0.2 volume of 10 M ammonium acetate and 1 volume of isopropanol, resulting the immediate formation of cloudy DNA precipitates. Precipitated DNA was then transferred by a pipet tip to a tube containing 75% (v/v) ethanol. This process was repeated twice to ensure complete removal of residual organic solvents. Afterwards, genomic DNA was dissolved in 1 mM Tris-HCl, pH 8.0, 0.1 mM EDTA by incubation at 50 °C for three hours.

### PCR amplification and next generation sequencing of sgRNAs

DNA fragments containing sgRNA sequences were amplified from isolated genomic DNA by two rounds of PCR using NEBNext Ultra II Q5 master mix (NEB). For the first round of PCR, forty-eight 50-μL PCR reactions (each containing 2 μg of genomic DNA template) were performed with the forward primer NGS-Lib-KO-Fwd-1 (5’-CCTACACGACGCTCTTCCGATCTNNNNNNNNNNNNNNNNNNGCTTTATATATCTTGTGGAAAGGACGAAACACC −3’) and the reverse primer NGS-Lib-KO-Rev-0 (5’-CAGACGTGTGCTCTTCCGATCTCCGACTCGGTGCCACTTTTTCAA-3’) with a thermal cycler program consisting of initial denaturation at 98°C for 5 minutes, followed by 16 cycles of (98°C denaturation for 10 seconds, 69°C annealing for 30 seconds, and 65°C extension for 45 seconds), and a final extension at 65°C for 5 minutes. Products of the first-round PCR were pooled and purified by a DNA clean and concentrator kit (Zymo) and diluted to 2 ng/μL. For the second round of PCR, six 50-μL PCR reactions (each containing 2 ng of the purified first-round PCR product) were performed for each sample with the forward primer NGS-Lib-KO-Fwd-2 (5’-AATGATACGGCGACCACCGAGATCTACACTCTTTCCCTACACGACGCTCTTCCGATCT-3’) and one of four indexed reverse primers (NGS-Lib-KO-Rev-1, 2, 3, 4) (5’-CAAGCAGAAGACGGCATACGAGAT[8-nucleotide index]GTGACTGGAGTTCAGACGTGTGCTCTTCCGATCT-3’). The same cycling conditions were used for second-round PCR as the first-round PCR except 18 cycles were used. Products of the second-round PCR reactions were subjected to electrophoresis on a 2% agarose gel. The expected ~300 bp amplicons were excised and extracted from the gel and sequenced by Illumina HiSeq PE150 (Novogene).

### Data analysis for genome-wide CRISPR knockout screening

CRISPR screening data were analyzed by MAGeCK (Model-based Analysis of Genome-wide CRISPR-Cas9 Knockout, v0.5.9.2)^19^ to discover candidate genes by comparing experimental condition (HQ461) with control condition (DMSO). Raw Fastq files were directly loaded into MAGeCK and the “count” command was used to collect read counts from Fastq files and to generate the sgRNA read count table. Next, the “test” command was used to perform statistical test from the count table, outputting log_2_ fold change, p value, and false discovery rate (FDR). The result table generated by MAGeCK was loaded into ggplot2^45^ in R to generate a volcano plot.

### Validation of top-ranking genes from CRISPR-Cas9 knockout screening

Two sgRNAs per gene were chosen from the Brunello library for DDB1 (5’-CATTGTCGATATGTGCGTGG-3’ and 5’-CTACCAACCTGCGATCACCA-3’), RBX1(5’-AGTACACTCTTCTGAAGTAG-3’ and 5’-ATGGATGTGGATACCCCGAG-3’), and UBE2G1 (5’-ACTTACTAAAGTGTATCTGG-3’ and 5’-ATGAAAAGCCAGAGGAACGC-3’). Annealed sgRNA oligos were cloned into the lentiCRISPR v2 vector. Lentiviral packaging was performed by co-transfecting the resulting plasmids with psPAX2 (Addgene 12260) and pMD2.G (Addgene 12259) into 293T cells. Media collected from transfected 293T cells was used to infect A549 cells at MOI ~5 for four days. The resulting cells were tested for their sensitivity to HQ461 using methods describe in the previous session cell viability assay.

### Isolation of HQ461 resistant HCT-116 clones for pooled whole exome-sequencing

Parental HCT-116 were plated sparsely on 10-cm plates to allow the isolation of ten individual clones. Each of these ten clones were expanded to one confluent 15-cm plate to establish ten independent cell populations. One million cells from each population were then plated on one 10-cm plate and treated with 20 μM HQ461 continuously for two weeks, with media change every 3 to 4 days. HQ461 resistant clones emerged from 5 out of the 10 populations. These clones were isolated, expanded, and tested for their sensitivity to HQ461 using cell viability assay. Five HQ461-resistant clones selected from independent populations were subjected to genomic DNA isolation as described in the previous session. Their genomic DNAs were then pooled in equal amounts for whole-exome sequencing at 200x average coverage (Novogene). As a control, genomic DNAs isolated from the corresponding cell populations before HQ461 selection were also pooled for whole-exome sequencing.

### Data analysis for whole-exome sequencing

Fastq files were aligned to the human reference genome version GRCh38 (hg38) using Burrows-Wheeler Aligner (BWA, v0.7.17)^46^ to generate sam files, which were then transformed to bam files through samtools (v1.10)^47^. Next, bam files were passed to the Genome Analysis Toolkit (GATK, v4.1.7)^22–24^ to call SNVs, deletions, and insertions. To call variants, several tools of GATK were utilized, including BaseRecalibrator to adjust base quality scores in the bam files using known variants in dbSNP database (build 146) and 1000 Genomes database, Mutect2 for variant calling, GetPileupSummaries and CalculateContamination to estimate and remove cross-sample contamination, CollectSequencingArtifactMetrics and FilterByOrientationBias to exclude sequence context-dependent artifacts, and FilterMutectCalls to filter out variants from some common sources of error. Finally, variants generated by GATK were annotated using snpEff (v4.5)^48^ and SnpSift (v4.3t)^49^.

### Sanger sequencing of *CDK12*

Genomic DNAs isolated from HQ461-resistant clones were used as template for amplification of a DNA sequence flanking glycine 731 of CDK12 with primers 5’-GTAAAACGACGGCCAGTGTTGTCCTCGTTATGGAGAAAGAA-3’ and 5’-CTGTGTCTTTGTCCTTGGCTTTAT-3’. PCR amplicons were Sanger sequenced with the M13F sequencing primer: 5’-GTAAAACGACGGCCAGT-3’.

### CRISPR-Cas9 knock-in of *CDK12* G731E or G731R alleles

An sgRNA targeting *CDK12* (5’-GTGGACAAGTTTGACATTAT-3’) was cloned into the pSpCas9(BB)-2A-eGFP (PX458) vector (Addgene 48138). For CDK12 G731E/R knock-in, 1 million A549 cells were nucleofected (using 4D-Nucleofector, Lonza) with PX458-sg*CDK12* and single-stranded oligodeoxynucleotides (G731E: 5’-ttatatacttggccataggttccttctccaataatTTCaataatgtcaaacttgtccacacagcgtttcccccagtc-3’; G731R: 5’-ttatatacttggccataggttccttctccaataatGCGaataatgtcaaacttgtccacacagcgtttcccccagtc-3’). Afterwards, cells were exposed to 5 μM 0461 for 2 weeks to select for cells with *CDK12* G731E/R knock-in. Cells that survived 0461 treatment were recovered and confirmed to have the correct G731E/R genomic conversion via Sanger sequencing.

### Generation of stable cell lines expressing 3xFLAG-CDK12

An ORF encoding *CDK12* isoform 2 (NM_015083.3) was PCR amplified from A549 cDNA and cloned into a pCDNA3.1-N-3xFLAG vector which was derived from the pCDNA3.1 vector. G731E and G731R mutations in *CDK12* were introduced by overlap extension PCR. After sequencing verification, wild-type and mutant 3xFLAG-*CDK12* sequences were subcloned into lentiCas9-Blast (Addgene 52962) by replacing the Cas9 ORF with 3xFLAG-*CDK12*. The resulting lenti-EF1α-3xFLAG-CDK12-P2A-BSD WT/GE/GR constructs were packaged into lentiviral vectors to transduce A549 cells. Stable cell lines were obtained by selection with 20 μg/ml of blasticidin.

### Antibodies for western blotting

Standard SDS-PAGE and western blotting procedures were used with the following modifications. For preparation of total lysates, cells were rinsed with DPBS to remove residual medium and then lysed in 20 mM HEPES-NaOH, pH 8.0, 10 mM NaCl, 2 mM MgCl_2_, 1% SDS freshly supplemented with 0.5 units/μL of benzonase and 1x cOmplete, Mini, EDTA-free protease inhibitor cocktail (Roche). Protein concentrations of the resulting lysates were quantified by the BCA method. Between 30 to 60 μg of proteins were resolved on SDS-PAGE and transferred to nitrocellulose membranes with a pore size of 0.5 μm.

Membranes were blocked in 5% nonfat milk PBST (0.1% v/v Tween-20) for 30 minutes before blotting with antibodies. The following primary antibodies were used by dilution in 5% nonfat milk PBST: anti-NRF2 (Abcam, ab62352, 1:4,000), anti-β-Actin-HRP (Huaxingbio, HX18271,1:10,000), anti-α-Tubulin-HRFP (MBL Life Science, PM054-7, 1:10,000), anti-DDB1 (Abcam, ab109027, 1:10,000), anti-RBX1 (Proteintech, 14895-1-AP, 1:5,000), anti-UBE2G1 (Proteintech, 12012-1-AP, 1:2,000), anti-CDK12 (Cell Signaling Technology, 11973S,1:4,000), anti-CCNK (Bethyl lab, A301-939A-T,1:4,000), anti-FLAG-HRP (Sigma, A8592, 1:10,000), anti-HA (BioLegend, 901533, 1:4,000), anti-RNA polymerase II subunit B1 (phospho CTD Ser-2) clone 3E10 (Merck, 04-1571-I, 1:5,000), and anti-RNA polymerase II CTD repeat YSPTSPS antibody clone 8WG16 (Abcam, ab817, 1:5,000). The following HRP-linked secondary antibodies were used by dilution in PBST: anti-rabbit IgG (Cell Signaling Technology, 7074S, 1:10,000), anti-mouse IgG (Zsbio, ZB-2305, 1:10,000), and anti-rat IgG, (Sino Biological, SSA005, 1:10,000). M5 HiPer ECL Western HRP Substrate (Mei5bio, MF074-01) was used for the detection of HRP enzymatic activity. Western blot images were taken with a VILBER FUSION FX7 imager.

### *In vivo* polyubiquitination of CCNK

Wild-type and G731E/R ORFs of 3×FLAG-*CDK12*, either full length or the kinase domain (from resides 715-1052), were cloned into the pCDNA3.1-N-3xFLAG vector. An ORF encoding CCNK (NM_001099402.2) was PCR amplified with A549 cDNA and cloned into a pCDNA3.1-C-3xHA vector. pCMV-8×His-Ub was a gift from Dr. William Kaelin at Dana Farber Cancer Institute of Harvard. To set up cells for the *in vivo* ubiquitination assay, 0.6 million HEK293T cells were seeded per well in 6-well tissue-culture plates. After overnight attachment, these cells were transfected with 500 ng of pCMV 8×His Ub, 25 ng of pCDNA3.1-CCNK-3xHA, and 50 ng of pCDNA3.1-3xFLAG-CDK12. Two days later, cells were pretreated with 1 μM of bortezomib for 2 hours, followed by treatment with DMSO or three doses of HQ461 (1, 3, 10 μM) for 4 hours. Cells from each well were rinsed once with DPBS and lysed in 500 μL of buffer 1 (25 mM Tris-HCl, pH 8.0, 8 M urea, 10 mM imidazole) freshly supplemented with 1 units/μL of benzonase and 1x cOmplete, Mini, EDTA-free protease inhibitor cocktail (Roche). Lysates were clarified by spinning at 15,000 xg at 4 °C for 10 minutes. Clarified lysate of each sample was mixed with 10 μL of Ni NTA magarose beads (SMART Life Scences, SM00801) prewashed with lysis buffer. Tubes containing the bead-lysate mixtures were rotated for 4 hours at room temperature followed by two washes with buffer 1, one wash with a 1:3 mixture of buffer 1: buffer 2 (25 mM Tris-HCl, pH 6.8, 20 mM imidazole), and one wash with buffer 2. His-Ub-conjugated proteins were eluted from the beads by boiling in 50 μL of 1x SDS sample buffer supplemented with 300 mM imidazole.

### Co-expression of 3xFLAG-CDK12 or 3xFLAG-CDK13 kinase domain with CCNK-3xHA

ORFs encoding residues 715-1052 of CDK12 and residues 693-1030 of CDK13 were cloned into a pCDNA3.1-N-3xFLAG vector. For setting up cell for co-transfection, 0.6 million 293T cells were seeded per well in 6-well tissue-culture plates and allowed to attach to plates overnight. These cells were then transfected with 50 ng of pCDNA3.1-3xFLAG-CDK12/13 kinase domain, 50 ng of pCDNA3.1-CCNK-3xHA, and 400 ng of the empty pCDNA3.1 vector. Twenty-four hours post transfection, cells were treated with HQ461 for 8 hours, and lysates were collected for western blotting with anti-FLAG and anti-HA antibodies.

### N-terminal 3xFLAG-tagging of endogenous *CDK12*

An sgRNA sequence (5’-GCCCAATTCAGAGAGACATG-3’) targeting the genomic region immediately downstream the CDK12 start codon was cloned into the PX458 vector (Addgene 48138). The 3xFLAG knock-in repair template was constructed in a pTOPO-TA vector (Mei5bio) containg a BSD-P2A-3xFLAG sequence flanked by two 500-bp homology arms matching upstream and downstream sequences of the *CDK12* genomic locus. For endogenous tagging of *CDK12*, 1 million A549 cells were nucleofected (using 4D-Nucleofector, Lonza) with 1μg of PX458-sg*CDK12* and 1 μg of the repair template. Selection with 30 μg/ml of blastistin was performed until clones appeared. Multiple clones were isolated and successful integration of N-terminal 3xFLAG tag was validated by western blotting with anti-FLAG-HRP (Sigma, A8592, 1:10,000).

### Co-immunoprecipitation of CCNK and DDB1 with 3xFLAG-CDK12 complex

Anti-FLAG M2 antibody (Sigma F3165) was coupled to magnetic epoxy beads (Beijing Yunci Technology Co.) at the ratio of 10 μg of anti-FLAG antibody/mg of beads in the presence of 1 M ammonium acetate and 0.1 M sodium phosphate, pH 7.4 at 37 °C overnight. A549 3xFLAG-*CDK12* knock-in cells were detached from plates by scraping, washed in DPBS, and then frozen in liquid nitrogen. Frozen cells were pulverized using a mixer mill MM 400 (Retsch) with two rounds of 1-minute ball milling at 30 Hz. Per experiment, 25 mg of grinded cell powder was solubilized with 250 μL of IP buffer (50 mM HEPES, pH 7.4, 300 mM NaCl, 0.1% Tween-20) supplemented with 1x cOmplete, Mini, EDTA-free protease inhibitor cocktail (Roche). The resulting lysates were centrifuged at 15,000 g for 10 minutes at 4 °C. Clarified lysates were supplemented with HQ461 or DMSO and incubated at 4°C for 30 minutes. Afterwards, 0.2 mg of anti-FLAG-conjugated magnetic beads were mixed with clarified lysates for 15 minutes on a rotating platform at 4 °C, followed by three washes with IP buffer supplemented with HQ461 or DMSO. Bound proteins were eluted from magnetic beads with 1 mg/ml of 3×FLAG peptide (Sigma F4799) with agitation at 4 °C for 30 min. Eluted proteins were subjected to SDS-PAGE and western blotting with anti-FLAG-HRP, anti-CCNK, and anti-DDB1.

### Expression and purification of ubiquitin and enzymes for*in vitro* ubiquitination assay

Human Ubiquitin ORF was cloned into pET28a vector with an N-terminal 6xHis tag. The resulting pET28a-6xHis-Ub plasmid was transformed into the *E. coli* strain BL21 (DE3) and grown at 37 °C until OD_600_ reached ~ 0.8. His-Ub expression was induced overnight by 0.5 mM IPTG at 18 °C. Cells were collected by centrifugation at 4000 g and resuspended with lysis buffer containing 50 mM Tris-HCl, pH 7.5, 500 mM NaCl, 20 mM imidazole, 5% glycerol. Cell lysis was performed by French Press. After 20,000 g centrifugation for 1 hour at 4 °C, the supernatant was incubated with Ni-NTA resin. After extensive wash with lysis buffer, His-Ub was eluted from Ni-NTA resin with 50 mM Tris-HCl, pH 7.5, 500 mM NaCl, 300 mM imidazole, 5% glycerol. Fractions containing His-Ub were pooled and concentrated, followed by gel filtration on a Superdex 200 10/300 GL column (GE Healthcare) in gel filtration buffer (50 mM Tris-HCl, pH7.5, 10 mM NaCl, 1 mM DTT). Fractions containing His-Ub were pooled and concentrated by a 3KDa MWCO Ultra centrifugal filter (Amicon), flash frozen with liquid nitrogen, and stored at −80℃ before use.

Human ORFs encoding UBA1 (residues 49-1058, NP_003325.2), UBE2G1 (NP_003333.1), UBE2D3 (NP_003331.1), CUL4A (NP_001008895.1) were cloned into pPB-CAG-1xFLAG vector. ORF encoding human RBX1 (NP_055063.1) was cloned into pPB-CAG vector without a tag. Plasmids were transfected into 293-F cells at the condition of 0.1 mg of plasmid for 100 million 293-F cells at a density at 1 million cells per ml. FLAG-UBA1, FLAG-UBE2G1, FLAG-UBE2D3 were separately expressed, whereas FLAG-CUL4A was co-expressed with RBX1. Cells were collected 48 hours post transfection by centrifugation at 1,000 g and then lysed in binding buffer (45 mM Tris-HCl, pH 7.8, 180 mM NaCl) supplemented with 0.2% Triton X-100, 1 mM DTT, and 1x cOmplete, Mini, EDTA-free protease inhibitor cocktail (Roche). After 20,000 g centrifugation, the clarified lysates were mixed with anti-FLAG M2 affinity gel (Sigma, A2220) for 2 hours at 4 °C on a rotator. After extensive washes with binding buffer plus 0.1% Triton X-100, FLAG-tagged proteins were eluted from beads by 0.2 mg/ml FLAG peptide (Sangon Biotech, T510060-0005). The protein-containing fractions were concentrated by ultracentrifugation and fractionated by an ENrich SEC650 gel filtration column (Bio-Rad) in binding buffer. Fractions containing target proteins were pooled, concentrated by ultrafiltration, flash frozen with liquid nitrogen and stored at −80°C.

Human DDB1(NP_001914.3) was cloned into pFastBac with an N-terminal Strep-tag and an 8×His tag. Expression and purification of His-DDB1 from Sf9 cells followed identical procedures as described in *Purification of DDB1∆BPB*.

### *In vitro* ubiquitination of CCNK

Per reaction, 3xFLAG-CDK12 in complex with CCNK was immunopurified from 1 million A549 cells stably expressing 3xFLAG-CDK12 (A549 lenti-3xFLAG-CDK12) using anti-FLAG antibody-conjugated magnetic beads. *In vitro* ubiquitination reactions were performed by mixing 3xFLAG-CDK12/CCNK captured on magnetic beads with 0.2 μM UBA1, 0.5 μM UBE2G1, 0.5 μM UBE2D3, 0.8 μM CUL4A-RBX1, 1 μM DDB1, and 100 μM ubiquitin in a buffer containing 50 mM HEPES, pH 7.5, 5 mM MgCl_2_, 5 mM ATP, 75 mM sodium citrate, and 0.1% Tween-20. HQ461 or DMSO were added to the reaction 30 minutes prior to the addition of enzymes. Reactions were incubated for 1 hour at 30 °C with agitation and then quenched with SDS sample buffer, followed by SDS-PAGE and western blotting with anti-CCNK antibody.

### Expression and purification of CDK12KD/CCNK∆C

Human CDK12KD (residues 715-1052) and human CCNK∆C (residues 11-267) were cloned into the baculoviral transfer vector pFastBac with an N-terminal 6xhis tag followed by a PreScission Protease cleavage site. G731E and G731R mutations were introduced to CDK12KD by overlap extension PCR. ORF of Cdk-activating kinase (CAK1) from *Saccharomyces cerevisiae* (Uniprot P43568) was cloned into pFastBac without a tag. Bacmid DNA was prepared in the *E. coli* strain DH10Bac and used to generate baculovirus by transfection into Sf9 cells. Sf9 or High Five cells were coinfected with baculovirus for CDK12KD (WT or G731E/R mutants), CCNK∆C, and yeast CAK1. Cells were lysed by sonication in binding buffer (50 mM HEPES, pH 7.5, 500 mM NaCl, 5% glycerol) supplemented with 20 mM imidazole and 1x cOmplete, Mini, EDTA-free protease inhibitor cocktail (Roche). The lysate was clarified by centrifugation at 15,000 g for 1 hour at 4°C. Recombinant proteins were purified by Ni NTA Beads (Smart Lifesciences, SA004010) with 30 mM imidazole for washing and 300 mM imidazole for elution. Fractions containing CDK12KD/CCNK∆C were treated with recombinant PreScission Protease at an enzyme to substrate ratio of 1:100 at 4 °C overnight to remove the 6xHis tag. The cleaved proteins were buffer-exchanged using 30 KDa MWCO ultra centrifugal filters (Amicon) to 50 mM MES, pH 6.5, 100 mM NaCl, 0.5 mM TCEP, and manually loaded onto a 5 mL HiTrap Q column (GE Healthcare). The flow-through containing CDK12KD/CCNK∆C was concentration by ultrafiltration, loaded onto ENrich SEC650 size exclusion column (Bio-Rad), and then fractionated in 50 mM HEPES, pH 7.5, 300 mM NaCl, 0.5 mM TCEP. Fractions containing CDK12KD/CCNK∆C were pooled and concentrated by ultrafiltration, flash frozen with liquid nitrogen, and stored at −80°C. For the AlphaScreen assay, His-tag removal was omitted from the above procedures.

### Purification of DDB1∆BPB

Human DDB1∆BPB (removing residues 400-704) was cloned into pFastBac with an N-terminal Strep-tag and an 8×His tag. Sf9 cells infected with DDB1∆BPB were lysed by sonication in binding buffer (50 mM Tris-HCl, pH 7.5, 500 mM NaCl, 10% glycerol, 2 mM TCEP, 1 mM PMSF, 10 mM imidazole) supplemented with 1x cOmplete, Mini, EDTA-free protease inhibitor cocktail (Roche). DDB1∆BPB was purified from clarified lysates by Ni NTA Beads and eluted in 50 mM Tris-HCl, pH 7.5, 500 mM NaCl, 10% glycerol, 2 mM TCEP, 1 mM PMSF, 300 mM imidazole. The fractions containing DDB1∆BPB were pooled, concentrated, and then subjected to gel filtration on an ENrich SEC650 size exclusion column (Bio-Rad) in 50 mM HEPES, pH 7.5, 300 mM NaCl, 0.5 mM TCEP.

### Purification of biotinylated FLAG-Avi-DDB1

Full-length human DDB1 was cloned into a pPB-CAG vector with an N-terminal FLAG-tag (for purification) and an Avi-tag (for biotinylation). The resulting pPB-CAG-FLAG-Avi-DDB1 construct was co-transfected with pPB-BirA (1:1 mass ratio) into 293-F cells at the condition of 0.1 mg of plasmid for 100 million 293-F cells at a density at 1 million cells per ml. D-Biotin (Targetmol, T1116) was added to the transfected cells at a final concentration of 50 μM. Cells were collected 48 hours post-transfection by 1,000 g centrifugation and resuspended in binding buffer (50 mM Tris-HCl, pH 7.5, 500 mM NaCl, 10% glycerol, 2 mM TCEP, 1 mM PMSF) supplemented with 1x cOmplete, Mini, EDTA-free protease inhibitor cocktail (Roche). After sonication, lysates were clarified by centrifugation at 15,000 g for 1 hour at 4°C. The supernatant containing recombinant proteins were incubated with anti-FLAG M2 affinity gel (Sigma, A2220) for 2 hours on a rotator at 4 °C. After extensive wash with binding buffer, FLAG-Avi-DDB1 was eluted from beads by 0.2 mg/ml FLAG peptide (Sangon Biotech, T510060-0005). The protein-containing fractions were concentrated by ultracentrifugation and further purified by gel filtration on an ENrich SEC650 size exclusion column (Bio-Rad).

### FLAG-DDB1:CDK12KD/CCNK∆C pull down assay

FLAG-Avi-DDB1 (18.2 μg) was mixed with CDK12KD/CCNK∆C (10 μg) at 1:1 molar ratio in 500 μL of pulldown assay buffer containing 50 mM HEPES, pH 7.5, 300 mM NaCl, 0.5 mM TCEP. The protein mix was then supplemented with DMSO or HQ461. After 30 minutes of incubation on ice, the protein-compound mix was incubated with anti-FLAG M2 affinity gel for another 30 minutes on a rotator at 4 °C. Protein-bound beads were washed three times with the pulldown assay buffer and bound proteins were eluted with 0.2 mg/mL FLAG peptide. Eluted proteins were analyzed by SDS-PAGE followed by coomassie blue staining.

### AlphaScreen assay

Biotinylated FLAG-Avi-DDB1 (3 μM) and 6xHis-CDK12KD/6xHis-CCNK∆C (WT or G731E/R mutants) (3 μM) were mixed in an assay buffer (50 mM HEPES, pH7.5, 300 mM NaCl, 0.5 mM TCEP, 0.1% NP-40, 0.1% BSA) and a serial dilution of HQ461.The protein-compound mix was transferred to 384-well plates (Corning 3572); 15 μL per well. After a 1-hour room temperature incubation, 10 μL of nickel chelate (Ni-NTA) acceptor beads (PerkinElmer, 6760619C) (1:100 diluted with assay buffer) was added per well. After incubation in the dark for 1 hour at room temperature, 10 μL of streptavidin donor beads (PerkinElmer, 6760619C) (1:100 diluted in assay buffer) was added per well. After another 1-hour incubation, AlphaScreen signal was measured on the EnVision multimode plate reader (PerkinElmer). For THZ531 competition, FLAG-Avi-DDB1, 6xHis-CDK12KD/6xHis-CCNK∆C mix was preincubated with 5 μM of THZ531 for 30 minutes on ice followed by the procedures described above.

### Chemical cross-linking mass spectrometry

A mixture of 23.4 μg of FLAG-Avi-DDB1 and 12.2 μg of CDK12KD/CCNK∆C (1:1 molar ratio) was incubated in the presence of 10 μM HQ461 for 30 minutes on ice, followed by cross-linking with 1 mM DSS (Thermo Fisher, A39267) or 1 mM BS3 (Thermo Fisher, A39266). The cross-linking reactions were incubated for 1 hour on a horizontal rotator at room temperature, followed by quenching with 20 mM ammonium bicarbonate for 20 minutes. Cross-linked samples were precipitated by 20% trichloroacetic acid (final concentration) for 30 minutes at −20 ºC. The resulting pellets were air-dried and then dissolved, assisted by sonication, in 8 M urea, 20 mM methylamine, 100 mM Tris-HCl, pH 8.5. After reduction (5 mM TCEP, room temperature, 20 minutes) and alkylation (10 mM iodoacetamide, room temperature, 15 minutes in the dark), the samples were diluted to 2 M urea with 100 mM Tris-HCl, pH 8.5. Denatured proteins were digested by trypsin at 1/50 (w/w) enzyme/substrate ratio at 37 °C overnight and the reactions were quenched with 5% formic acid (final concentration).

### Mass spectrometry analysis of cross-linked DDB1/CDK12KD/CCNK∆C

The LC-MS/MS analysis was performed on an Easy-nLC 1000 HPLC (Thermo Fisher Scientific) coupled to a Q-Exactive HF mass spectrometer (Thermo Fisher Scientific). Peptides were loaded on a pre-column (75 μm ID, 4 cm long, packed with ODS-AQ 12 nm-10 μm beads) and separated on an analytical column (75 μm ID, 12 cm long, packed with Luna C18 1.9 μm 100 Å resin) with a 60 min or 120 min linear gradient at a flow rate of 250 nl/min. The 60 min linear gradient was as follows: 0-5% B in 1 min, 5-30% B in 45 min, 30-100% in 4 min, 100% for 10 min (A = 0.1% FA, B = 100% ACN, 0.1% FA); the 120 min linear gradient was as follows: 0-5% B in 1 min, 5-30% B in 100 min, 30-100% in 4 min, 100% for 15 min (A = 0.1% FA, B = 100% ACN, 0.1% FA). The top fifteen most intense precursor ions from each full scan (resolution 60,000) were isolated for HCD MS2 (resolution 15,000; NCE 27) with a dynamic exclusion time of 30 s. Precursors with 1+, 2+, more than 7+ or unassigned charge states were excluded.

### Identification of cross-linked peptides using pLink2

The pLink2^36^ search parameters were as follows: instrument, HCD; precursor mass tolerance, 20 ppm; fragment mass tolerance 20 ppm; cross-linker, BS3/DSS (cross-linking sites K and protein N terminus, cross-link mass-shift 138.068, mono-link mass-shift 156.079); peptide length, minimum 6 amino acids and maximum 60 amino acids per chain; peptide mass, minimum 600 and maximum 6,000 Da per chain; enzyme, Trypsin, with up to three missed cleavage sites per chain; Carbamidomethyl[C], Oxidation[M], Deamidated[N] and Deamidated[Q] as variable modification. The results were filtered by requiring FDR < 1%, E-value <0.0001, spectra count > 2.

### Differential Scanning Fluorimetry (DSF)

CDK12KD/CCNK∆C (WT or G731E/R) and DDB1∆BPB were diluted to 5 μM with DSF assay buffer (50 mM HEPES pH 7.5, 300 mM NaCl, 0.5 mM TCEP). 50 μM HQ461 or 10 μM HBx was added to diluted proteins (final DMSO concentration 0.25%). Twenty μL of protein-compound mix was aliquoted per qPCR tube (Labtidebiotech, P01-0803E) followed by the addition of 5 μL of 1:400 diluted SYPRO Orange (Sigma, S5692). After incubation on ice for 20 minutes, fluorescence measurements were performed using a Bio-Rad CFX96 Realtime PCR instrument. During the DSF experiment, the temperature was increased from 10 to 95°C at an increment of 0.5°C with equilibration time of 10 seconds at each temperature prior to measurement. Tm was defined as the temperature corresponding to the maximum value of the first derivative of fluorescence transition.

### CCNK-luc degradation assay

The ORF for CCNK was fused with the firefly luciferase sequence and inserted into the pCDNA3.1 vector. For setting up cell for co-transfection, 0.6 million 293T cells were seeded per well in 6-well plates and allowed to attach to plates overnight. These cells were then transfected with 50 ng of pCDNA3.1-3xFLAG-CDK12 kinase domain, 50 ng of pCDNA3.1-CCNK-luc, and 400 ng of the empty pCDNA3.1 vector. Twenty-four hours later, transfected cells were dissociated by trypsin digestion and then plated in 96-well white plates. After overnight attachments, cells were then treated with HQ461 and its analog derivatives for eight hours followed by measurements of luciferase activity with Bright-Glo luciferase assay system (Promega). Half maximal CCNK-degradation concentrations (DC_50_) of HQ461 and its active analogs were interpolated by curve fitting with GraphPad Prism using baseline correction (by normalizing to DMSO control), the asymmetric (five parameter) equation, and least squares fit.

### RNA isolation and RT-qPCR

Total RNA was extracted from HQ461-treated A549 cells using TRNzol (Tiangen, DP405). Reverse transcription of mRNA into cDNA was performed using the 5X All-In-One RT mastermix kit (Abm,G488) with AccuRT genomic DNA removal kit (Abm, G489), and qPCR was performed using TB Green Premix Ex Taq (Tli RNaseH Plus) (Takara, RR420) with the CFX96 Real-Time PCR Detection System (Bio-Rad). GAPDH was used for normalization. Primer sequences are as follows: GAPDH-forward: 5’-GCTCTCTGCTCCTCCTGTTC-3’; GAPDH-reverse: 5’-ACGACCAAATCCGTTGACTC-3’; BRCA1-forward: 5’-GGCTGTGGGGTTTCTCAGAT-3’; BRCA1-reverse: 5’-TTCATGGAGCAGAACTGGTG-3’; BRCA2-forward: 5’-TTCATGGAGCAGAACTGGTG-3’; BRCA2-reverse: 5’-AGGAAAAGGTCTAGGGTCAGG-3’; ATR-forward: 5’-CGCTGAACTGTACGTGGAAA-3’; ATR-reverse: 5’-CAATAAGTGCCTGGTGAACATC-3’; ERCC4-forward: 5’-TAGACCTAGTAAGAGGCACAG-3’; ERCC4-reverse: 5’-GAGTGAGGATGTAATCTCCAA-3’. Each experiment was performed with three technical replicates.

### Colony formation assay after CCNK depletion by CRISPR-Cas9

An sgRNA sequence (5’-GTGAACCGAGCGCCCTCTCGG-3’) targeting *CCNK* was cloned into lentiCRISPR v2. An sgRNA-resistant ORF of *CCNK* (by replacing the sgRNA target sequence to 5’-taTcgTcgTgaAggTgcGcgCttTa-3’) was cloned into lentiCas9-blast by replacing Cas9 ORF with the sgRNA-resistant *CCNK* ORF. The resulting plasmid was packaged into lentiviral vector to stably infect A549 cells. Parental A549 cells or A549 cells expressing sgRNA-resistant *CCNK* were infected with high MOI with lentivirus carrying sg*NTC* or sg*CCNK*. One day post infection, 1,000 cells were plated per well in a 6-well tissue-culture plate. Cells were grown for 10 days with medium change every 2 to 3 days. The resulting cell colonies were fixed in 4% PFA in PBS, stained with crystal violet (Beyotime, C0121) for 20 minutes at room temperature, and then destained with water. Images were taken with VILBER FX7 imager.

### Preparation analogues of HQ461

All reactions were carried out under an atmosphere of nitrogen in flame-dried glassware with magnetic stirring unless otherwise indicated. Commercially obtained reagents were used as received. Solvents were dried by passage through an activated alumina column under argon. Liquids and solutions were transferred via syringe. All reactions were monitored by thin-layer chromatography with E. Merck silica gel 60 F254 pre-coated plates (0.25 mm). Structures of the target compounds in this work were assigned by use of NMR spectroscopy and MS spectrometry. The purities of all compounds were >95% as determined on Waters HPLC with 2998PDA and 3100MS detectors, using ESI as ionization. Pre-HPLC is used to separate and refine high-purity target compounds. ^1^H and ^13^C NMR spectra were recorded on Varian Inova-400 spectrometers. Data for 1H NMR spectra are reported relative to CDCl3 (7.26 ppm), CD3OD (3.31 ppm), or DMSO-d6 (2.50 ppm) as an internal standard and are reported as follows: chemical shift (δ ppm), multiplicity (s = singlet, d = doublet, t = triplet, q = quartet, sept = septet, m = multiplet, br = broad), coupling constant J (Hz), and integration.

## General procedure A

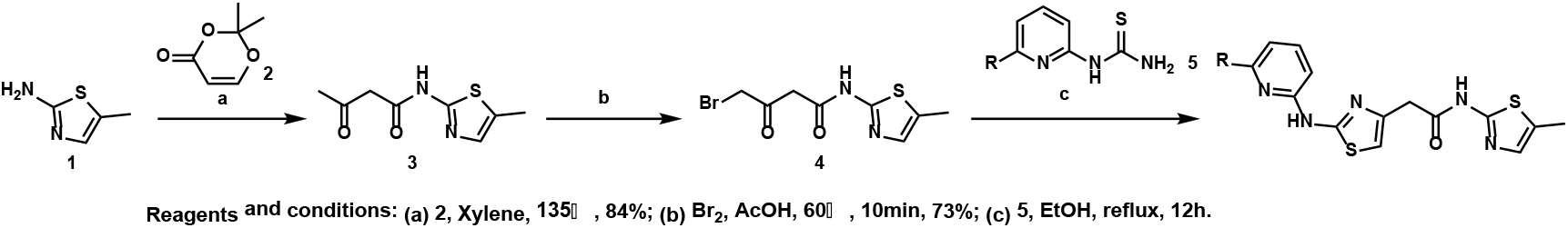

### Supplementary Scheme 1: General procedure A

#### Synthesis of N-(5-methylthiazol-2-yl)-3-oxobutanamide (3)

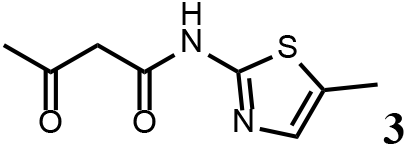

To a solution of 5-methylthiazol-2-amine (1) (0.5 g; 4.39 mmol) in xylene (10 ml) was added 2, 2-dimethyl-4H-1, 3-dioxin-4-one (2) (685 mg; 4.83 mmol). The mixture was stirred for 12 h at 135℃, then cooled to room temperature. The solution was filtered and washed with petroleum ether, after filtration, the solid was used directly without purification (3) (0.73 g; 3.69 mmol, 84%).

^1^H NMR (400 MHz, DMSO-d6) δ 11.95 (s, 1H), 7.13 (s, 1H), 3.66 (s, 2H), 2.34 (s, 3H), 2.19 (s, 3H); ^13^C NMR (101 MHz, DMSO-d6) δ 202.68, 165.26, 156.30, 135.18, 126.70, 51.22, 30.70, 11.53; LC-MS (*m/z*): [M+H]^+^ calculated for C_8_H_10_N_2_O_2_S, 199.0463; found, 198.8507.

#### Synthesis of 4-bromo-N-(5-methylthiazol-2-yl)-3-oxobutanamide (4)

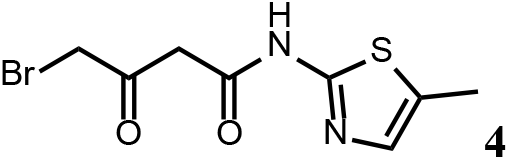

To a solution of N-(5-methylthiazol-2-yl)-3-oxobutanamide (3) (2.0 g; 10.1 mmol) in AcOH (30 ml) was added Br_2_ (2.0 g; 11.1 mmol) at 60℃. The mixture was stirred for 10min at 60℃, then cooled to room temperature. The reaction mixture was quenched with saturated aqueous Na_2_SO_3_ solution, extracted with EtOAc and the organic layer dried over Na_2_SO_4_, filtered and evaporated. The product was purified by column chromatography on silica gel to obtain 4-bromo-N-(5-methylthiazol-2-yl)-3-oxobutan amide (4) (2.0 g; 7.25 mmol, 73%).

^1^H NMR (400 MHz, DMSO-d_6_) δ 12.02 (s, 1H), 7.14 (s, 1H), 4.48 (s, 2H), 3.83 (s, 2H), 2.34 (s, 3H). ^13^C NMR (101 MHz, DMSO-d_6_) δ 196.04, 164.67, 156.24, 135.15, 126.89, 47.91, 37.59, 11.55; LC-MS (*m/z*): [M+H]^+^ calculated for C_8_H_9_BrN_2_O_2_S, 276.9568; found, 277.1675.

#### Synthesis of the HQ461 and analogues-A

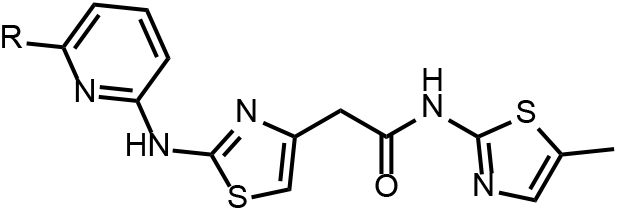

To a solution of above 4 (10 mg; 0.036 mmol) in EtOH (2 ml) was added the pyridinyl thiourea 5 (0.054 mmol). The mixture was stirred for 12 h at 70℃, then cooled to room temperature. The mixture was evaporated to dryness and purified by silica gel chromatography to get HQ461 analogues as HBr salt.

#### 2-(2-((6-methylpyridin-2-yl)amino)thiazol-4-yl)-N-(5-methylthiazol-2-yl)acetamide (HQ461)

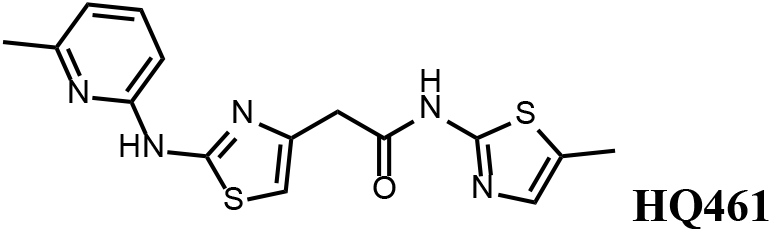

Pale yellow solid; Yield 52%; ^1^H NMR (400 MHz, DMSO-d6) δ 12.12 (s, 1H), 11.67 (s, 1H), 7.66 (t, J = 7.7 Hz, 1H), 7.14 (d, J = 1.2 Hz, 1H), 6.88 (t, J = 10.2 Hz, 3H), 3.85 (s, 2H), 2.47 (s, 3H), 2.33 (s, 3H); ^13^C NMR (101 MHz, DMSO-d_6_) δ 167.50, 161.13, 156.59, 154.46, 149.70, 140.66, 138.69, 134.87, 126.86, 117.62, 110.42, 109.70, 36.39, 22.99, 11.61; LC-MS (*m/z*): [M+H]^+^ calculated for C_15_H_15_N_5_OS_2_, 346.0718; found, 346.3456.

#### 2-(2-((6-fluoropyridin-2-yl)amino)thiazol-4-yl)-N-(5-methylthiazol-2-yl)acetamide (HQ001)

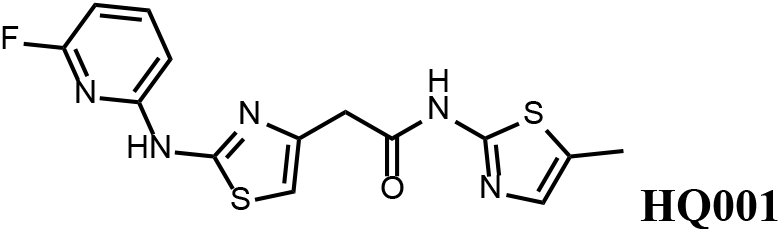

Pale yellow solid; Yield 50%; ^1^H NMR (400 MHz, CDCl_3_: MeOD= 4:1) δ 7.53 (dd, J = 15.9, 7.9 Hz, 1H), 6.87 (s, 1H), 6.60 (d, J = 7.7 Hz, 1H), 6.49 (s, 1H), 6.28 (d, J = 7.8 Hz, 1H), 3.61 (s, 2H), 2.20 (s, 3H). ^19^F NMR (376 MHz, CDCl3:MeOD= 4:1) δ −66.70; LC-MS (*m/z*): [M+H]^+^ calculated for C_14_H_12_FN_5_OS_2_, 350.0467; found, 350.0437.

#### 2-(2-((6-chloropyridin-2-yl)amino)thiazol-4-yl)-N-(5-methylthiazol-2-yl)acetamide (HQ002)

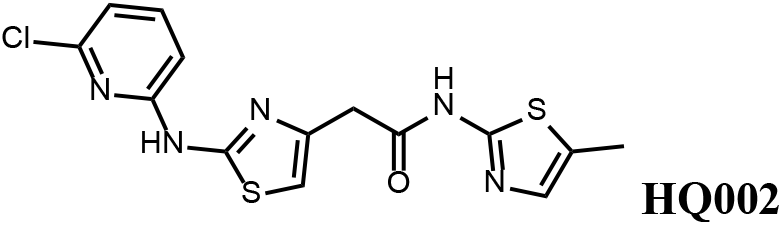

Pale yellow solid; Yield 35%; ^1^H NMR (400 MHz, CDCl_3_: MeOD= 4:1) δ 7.52 (dt, J = 11.0, 7.6 Hz, 1H), 7.01 (s, 1H), 6.89 – 6.82 (t, J=7.2 Hz, 1H), 6.78 (dd, J = 8.2, 3.2 Hz, 1H), 6.58 (s, 1H), 3.76 (s, 2H), 2.35 (s, 3H). LC-MS (*m/z*): [M+H]^+^ calculated for C_14_H_12_ClN_5_OS_2_, 366.0172; found, 366.2866.

#### 2-(2-((6-bromopyridin-2-yl)amino)thiazol-4-yl)-N-(5-methylthiazol-2-yl)acetamide (HQ003)

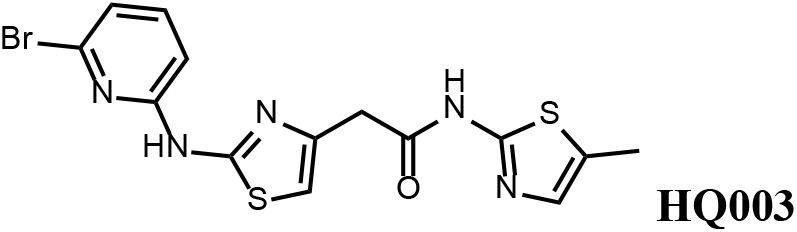

Pale yellow solid; Yield 53%; ^1^H NMR (400 MHz, CDCl_3_: MeOD= 4:1) δ 7.41 (t, J = 7.9 Hz, 1H), 6.99 (m, 2H), 6.79 (d, J = 8.3 Hz, 1H), 6.58 (s, 1H), 3.74 (s, 2H), 2.32 (s, 3H); LC-MS (*m/z*): [M+H]^+^ calculated for C_14_H_12_BrN_5_OS_2_, 409.9667; found, 410.2289.

#### 2-(2-((6-ethylpyridin-2-yl)amino)thiazol-4-yl)-N-(5-methylthiazol-2-yl)acetamide (HQ004)

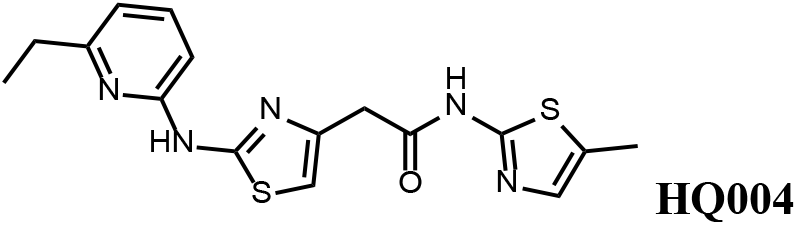

Pale yellow solid; Yield 53%; ^1^H NMR (400 MHz, CDCl3: MeOD= 4:1) δ 7.52 (t, J = 7.8 Hz, 1H), 7.06 (s, 1H), 6.76 (d, J = 7.6 Hz, 1H), 6.64 (d, J = 7.8 Hz, 1H), 6.59 (s, 1H), 3.79 (s, 2H), 2.83 (q, J = 7.2, 14.8 Hz, 2H), 2.39 (s, 3H), 1.38 (t, J = 7.2, 3H); LC-MS (*m/z*): [M+H]^+^ calculated for C_16_H_17_N_5_OS_2_, 360.0875; found, 360.0505.

#### 2-(2-((6-(hydroxymethyl)pyridin-2-yl)amino)thiazol-4-yl)-N-(5-methylthiazol-2-yl)acetamide (HQ005)

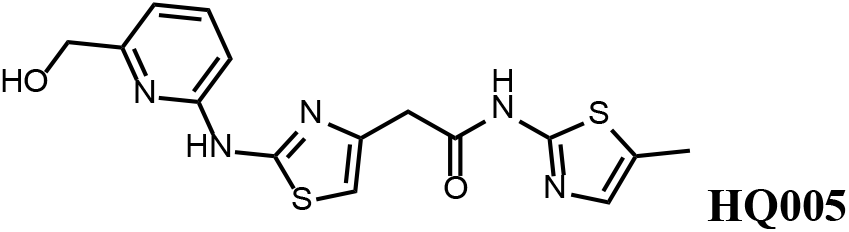

Pale yellow solid; Yield 33%; ^1^H NMR (400 MHz, CDCl3) δ 11.93 (s, 1H), 9.83 (s, 1H), 7.61 (t, J = 7.8 Hz, 1H), 7.08 (s, 1H), 6.86 (d, J = 7.6 Hz, 1H), 6.72 (d, J = 7.7 Hz, 1H), 6.63 (s, 1H), 4.80 (s, 2H), 3.80 (s, 2H), 2.40 (s, 3H); LC-MS (*m/z*): [M+H]^+^ calculated for C_15_H_15_N_5_O_2_S_2_, 362.0667; found, 362.2984.

#### Tert-butyl(6-((4-(2-((5-methylthiazol-2-yl)amino)-2-oxoethyl)thiazol-2-yl)amino)pyridin-2-yl)carbamate (HQ007-01)

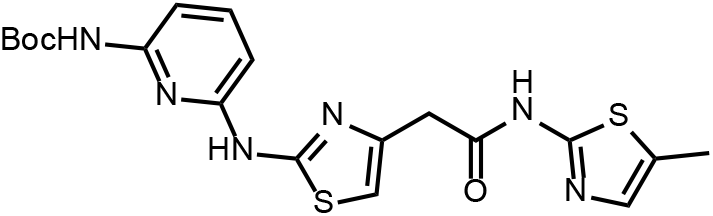

White solid; Yield 35%; ^1^H NMR (400 MHz, CDCl_3_: MeOD= 4:1) δ 7.60 (t, J = 7.6 Hz, 1H), 7.50 (d, J = 7.7 Hz, 1H), 7.06 (s, 1H), 6.56 (s, 1H), 6.51 (d, J = 7.9 Hz, 1H), 3.79 (s, 2H), 2.39 (s, 3H), 1.549 (s, 9H); LC-MS (*m/z*): [M+H]^+^ calculated for C_19_H_22_N_6_O_3_S_2_, 447.1195; found, 447.4275.

#### 2-(2-((6-aminopyridin-2-yl)amino)thiazol-4-yl)-N-(5-methylthiazol-2-yl)acetamide (HQ007)

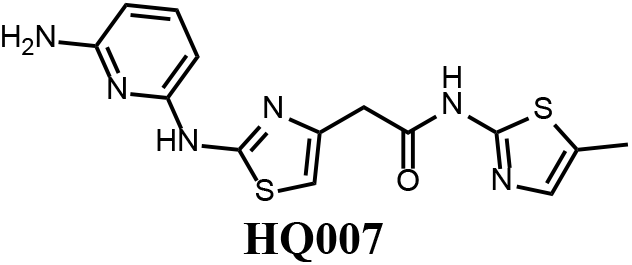

To a solution of HQ007-01 (3 mg) in dichloromethane (1 ml) was added Trifluoroacetic acid (0.5 ml). The mixture was stirred for 2 h at room temperature. The mixture was evaporated to dryness and purified by silica gel chromatography to get 2-(2-((6-aminopyridin-2-yl) amino)thiazol-4-yl)-N-(5-methylthiazol-2-yl)acetamide as TFA salt (2.4 mg, 80%).

^1^H NMR (400 MHz, CDCl3: MeOD= 4:1) δ 7.42 (t, J = 8.3 Hz, 1H), 6.83 (s, 1H), 6.66 (s, 1H), 6.13 (d, J = 7.9 Hz, 1H), 6.06 (d, J = 8.1 Hz, 1H), 3.65 (s, 2H), 2.13 (s, 3H); LC-MS (*m/z*): [M+H]^+^ calculated for C_14_H_14_N_6_OS_2_, 347.0671; found, 347.2157.

#### N-(5-methylthiazol-2-yl)-2-(2-((6-(trifluoromethyl)pyridin-2-yl)amino)thiazol-4-yl)acetamide (HQ009)

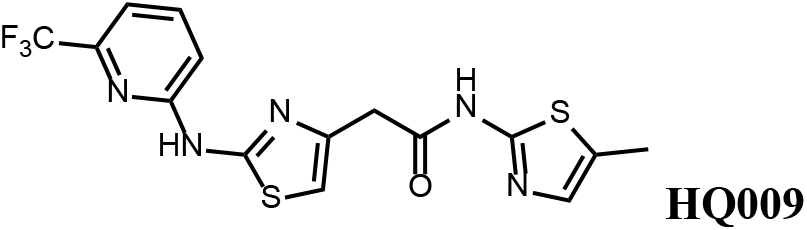

White solid; Yield 52%; ^1^H NMR (400 MHz, CDCl3: MeOD= 4:1) δ 7.71 (t, J = 8.1 Hz, 1H), 7.18 (d, J = 7.6 Hz, 1H), 7.03 (d, J = 8.1 Hz, 1H), 6.99 (s, 1H), 6.59 (s, 1H), 3.75 (s, 2H), 2.33 (s, 3H). ^19^F NMR (376 MHz, CDCl3:MeOD= 4:1) δ −68.59; LC-MS (*m/z*): [M+H]^+^ calculated for C_15_H_12_F_3_N_5_OS_2_, 400.0435; found, 400.3673.

#### N-(5-methylthiazol-2-yl)-2-(2-(pyridin-2-ylamino)thiazol-4-yl)acetamide (HQ010)

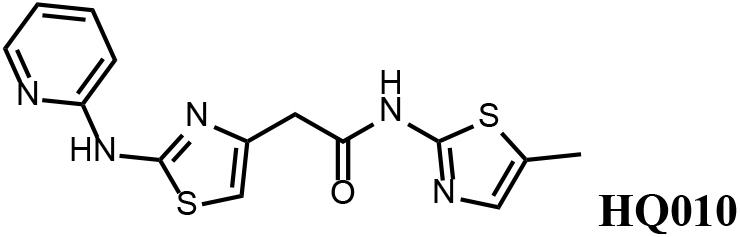

Pale yellow solid; Yield 53%; ^1^H NMR (400 MHz, CDCl3: MeOD= 4:1) δ 8.28 (d, J = 4.6 Hz, 1H), 7.57 (t, J=7.4Hz, 1H), 6.99 (s, 1H), 6.85 (m, 2H), 6.51 (s, 1H), 3.73 (s, 2H), 2.33 (s, 3H); LC-MS (*m/z*): [M+H]^+^ calculated for C_14_H_13_N_5_OS_2_, 332.0562; found, 332.3505.

## General procedure B

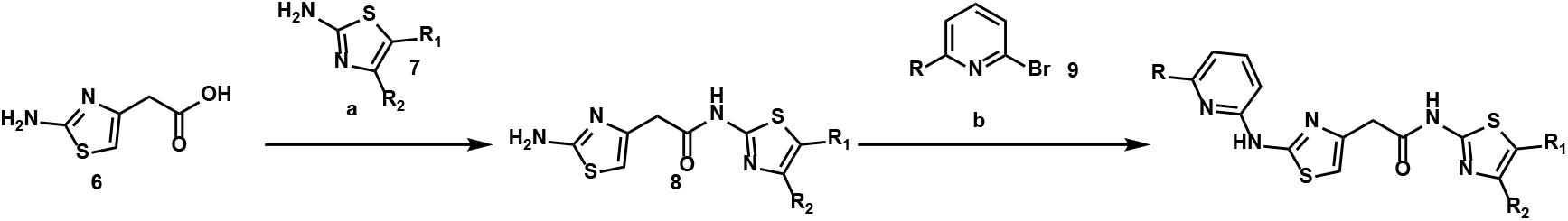

### Supplementary Scheme 2: General procedure B

#### Synthesis of 2-(2-aminothiazol-4-yl)-N-(5-methylthiazol-2-yl)acetamide (HQ011)

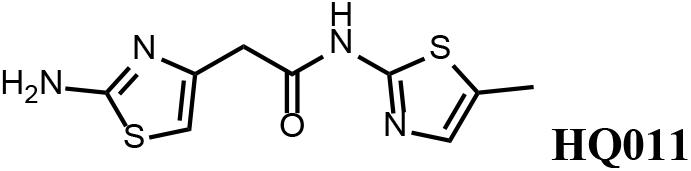

To a mixture of 2-(2-aminothiazol-4-yl)acetic acid (6) (1.00 g; 6.33 mmol) and 5-methylthiazol-2-amine (1) (722 mg; 6.33 mmol) in dimethylformamide (15 ml) was added 1-Ethyl-3-(3-dimethylaminopropyl) carbodiimide hydrochloride (1.82 g; 9.50 mmol) and dimethylaminopyridine (154mg; 1.27 mmol) under argon atmosphere. The mixture stirred at room temperature for 3 hours. The reaction mixture was quenched with water, extracted with EtOAc and the organic layer dried over Na_2_SO_4_, filtered and evaporated. The product was purified by column chromatography on silica gel to obtain 2-(2-aminothiazol-4-yl)-N-(5-methylthiazol-2-yl) acetamide (**7**) (1.15 g; 4.53 mmol, 72%).

^1^H NMR (400 MHz, DMSO-d6) δ 11.97 (s, 1H), 7.12 (s, , 1H), 6.95 (s, 2H), 6.32 (s, 1H), 3.56 (s, 2H), 2.33 (s, 3H); ^13^C NMR (101 MHz, DMSO-d6) δ 168.82, 168.27, 156.59, 145.33, 135.17, 126.54, 103.53, 38.52, 11.57; LC-MS (*m/z*): [M+H]^+^ calculated for C_9_H_10_N_4_OS_2_, 255.0296; found, 255.2680.

#### 2-(2-aminothiazol-4-yl)-N-(4-(furan-2-yl)thiazol-2-yl)acetamide (HQ013)

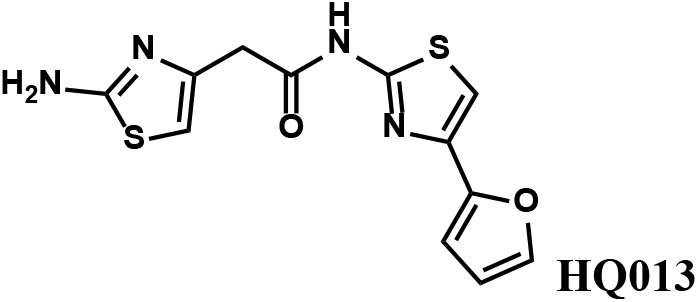

To a mixture of 2-(2-aminothiazol-4-yl)acetic acid (6) (100 mg; 0.63 mmol) and 4-(furan-2-yl)thiazol-2-amine (105 mg; 0.63 mmol) in dimethylformamide (5 ml) was added 1-Ethyl-3-(3-dimethylaminopropyl) carbodiimide hydrochloride (182 mg; 0.95 mmol) and dimethylaminopyridine (15 mg; 0.13 mmol) under argon atmosphere. The mixture stirred at room temperature for 3 hours. The reaction mixture was quenched with water, extracted with EtOAc and the organic layer dried over Na_2_SO_4_, filtered and evaporated. The product was purified by column chromatography on silica gel to obtain 2-(2-aminothiazol-4-yl)-N-(4-(furan-2-yl)thiazol-2-yl)acetamide (HQ013) (124 mg; 0.41 mmol, 65%).

^1^H NMR (400 MHz, DMSO-d6) δ 12.43 (s, 1H), 7.72 (dd, J = 1.8, 0.8 Hz, 1H), 7.32 (s, 1H), 6.93 (s, 2H), 6.67 (d, J = 2.7 Hz, 1H), 6.58 (dd, J = 3.3, 1.8 Hz, 1H), 6.35 (s, 1H), 3.60 (s, 2H); LC-MS (*m/z*): [M+H]^+^ calculated for C_12_H_10_N_4_O_2_S_2_, 307.0245; found, 307.1883.

#### Synthesis of the HQ461 and analogues-B

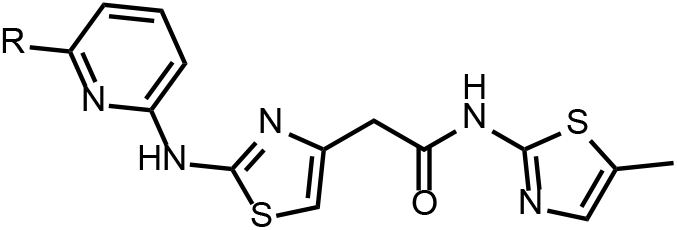

A mixture of HQ011 (20mg, 0.08mmol), 9 (0.096 mmol), Xantphos (5.6 mg, 0.0096 mmol) Tris(dibenzylideneacetone)dipalladium (8.8 mg, 0.0096 mmol), tBuOK (21 mg, 0.19 mmol) in 1,4-dioxane(2.0 ml) under argon atmosphere was stirred at 110 °C for 12 h. The mixture was filtered and concentrated in vacuum and the crude product purified by column chromatography on silica gel.

#### 2-(2-((6-methoxypyridin-2-yl)amino)thiazol-4-yl)-N-(5-methylthiazol-2-yl)acetamide (HQ006)

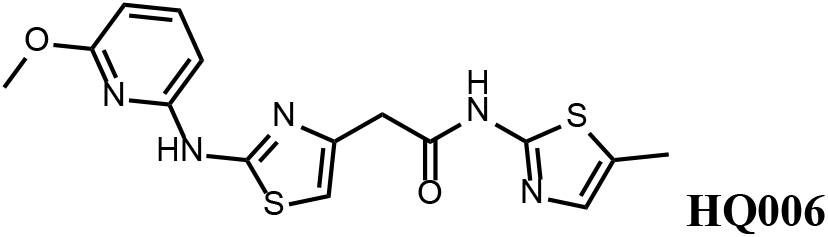

Pale yellow solid; Yield 33%; ^1^H NMR (400 MHz, CDCl_3_) δ 11.60 (s, 1H), 9.36 (s, 1H), 7.51 (t, J = 7.9 Hz, 1H), 7.09 (s, 1H), 6.62 (s, 1H), 6.35 (dd, J = 7.9, 3.3 Hz, 2H), 4.08 (s, 3H), 3.79 (s, 2H), 2.40 (s, 3H); LC-MS (*m/z*): [M+H]^+^ calculated for C_15_H_15_N_5_O_2_S_2_, 362.0667; found, 362.3709.

#### 2-(2-((6-(1H-pyrazol-1-yl)pyridin-2-yl)amino)thiazol-4-yl)-N-(5-methylthiazol-2-yl)acetamide (HQ008)

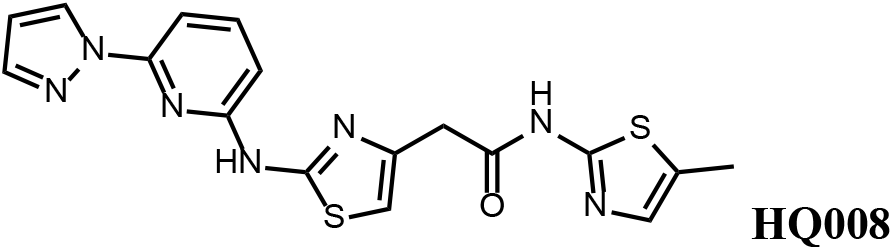

Pale yellow solid; Yield 25%; ^1^H NMR (400 MHz, DMSO-d6) δ 12.09 (s, 1H), 11.63 (s, 1H), 8.75 (d, J = 2.2 Hz, 1H), 7.91 – 7.81 (m, 2H), 7.42 (d, J = 7.9 Hz, 1H), 7.14 (s, 1H), 6.94 – 6.88 (m, 2H), 6.64 (S, 1H), 3.80 (s, 2H), 2.33 (s, 3H); LC-MS (*m/z*): [M+H]^+^ calculated for C_17_H_15_N_7_OS_2_, 398.0779; found, 398.3370.

#### N-(4-(furan-2-yl)thiazol-2-yl)-2-(2-((6-methylpyridin-2-yl)amino)thiazol-4-yl)acetamide (HQ012)

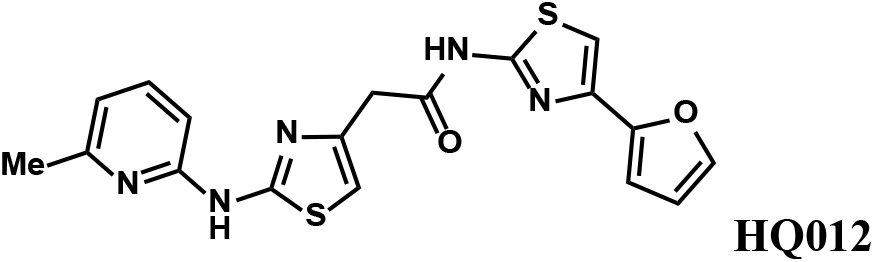

A mixture of HQ013 (20mg, 0.08mmol), 2-bromo-6-methylpyridine (16.4 mg, 0.096 mmol), Xantphos (5.6 mg, 0.0096 mmol), Tris(dibenzylideneacetone)dipalladium (8.8 mg, 0.0096mmol), tBuOK (21 mg, 0.19 mmol) in 1,4-dioxane (2.0 ml) under argon atmosphere was stirred at 110 °C for 12 h. The mixture was filtered and concentrated in vacuum and the crude product purified by column chromatography on silica gel to obtain N-(4-(furan-2-yl)thiazol-2-yl)-2-(2-((6-methylpyridin-2-yl)amino)thiazol-4-yl) acetamide (HQ012) (9.8 mg; 0.026 mmol, 21%).

^1^H NMR (400 MHz, CDCl_3_) δ 11.99 (s, 1H), 9.40 (s, 1H), 7.54 – 7.47 (m, 1H), 7.43 (d, J = 1.8 Hz, 1H), 7.08 (s, 1H), 6.76 (d, J = 7.4 Hz, 1H), 6.69 (d, J = 3.3 Hz, 1H), 6.64 (d, J = 8.1 Hz, 1H), 6.57 (s, 1H), 6.48 – 6.43 (m, 1H), 3.82 (s, 2H), 2.55 (d, J = 2.9 Hz, 3H); LC-MS (*m/z*): [M+H]^+^ calculated for C_18_H_15_N_5_O_2_S_2_, 398.0667; found, 398.3370.

#### Synthesis of 6-bromo-N-(4-(2-((5-methylthiazol-2-yl)amino)-2-oxoethyl)thiazol-2-yl) Picolinamide (HQ014)

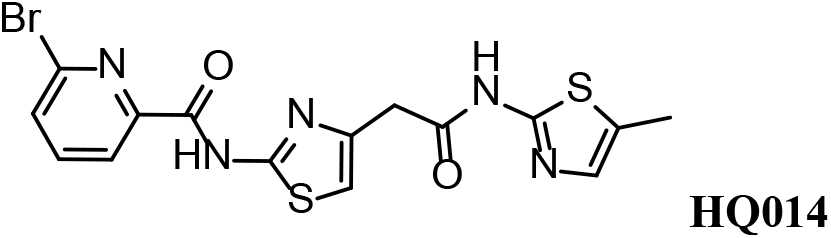

A mixture of HQ011 (25mg, 0.10mmol), 6-bromopicolinic acid (20mg, 0.10 mmol), 1-Ethyl-3-(3-dimethylaminopropyl) carbodiimide hydrochloride (28mg, 0.15mmol) dimethylaminopyridine (1.22 mg, 0.01mmol) in dimethylformamide (2.0 ml) under argon atmosphere was stirred at room temperature for 12 h. The reaction mixture was quenched with water, extracted with EtOAc and the organic layer dried over Na_2_SO_4_, filtered and evaporated. The product was purified by column chromatography on silica gel to obtain HQ014 (28 mg; 0.064 mmol, 83%).

^1^H NMR (400 MHz, CDCl_3_: MeOD= 4:1) δ 8.18 (dd, J = 7.5, 0.6 Hz, 1H), 7.76 (t, J = 7.7 Hz, 1H), 7.71 – 7.66 (m, 1H), 6.99 (s, 1H), 6.87 (s, 1H), 3.82 (s, 2H), 2.32 (s, 3H); LC-MS (*m/z*): [M+H]^+^ calculated for C_15_H_12_BrN_5_O_2_S_2_, 437.9616; found, 438.2910.

